## Supplemental file for "Ligand Response of Guanidine-IV riboswitch at Single-molecule Level"

### Supplemental Information

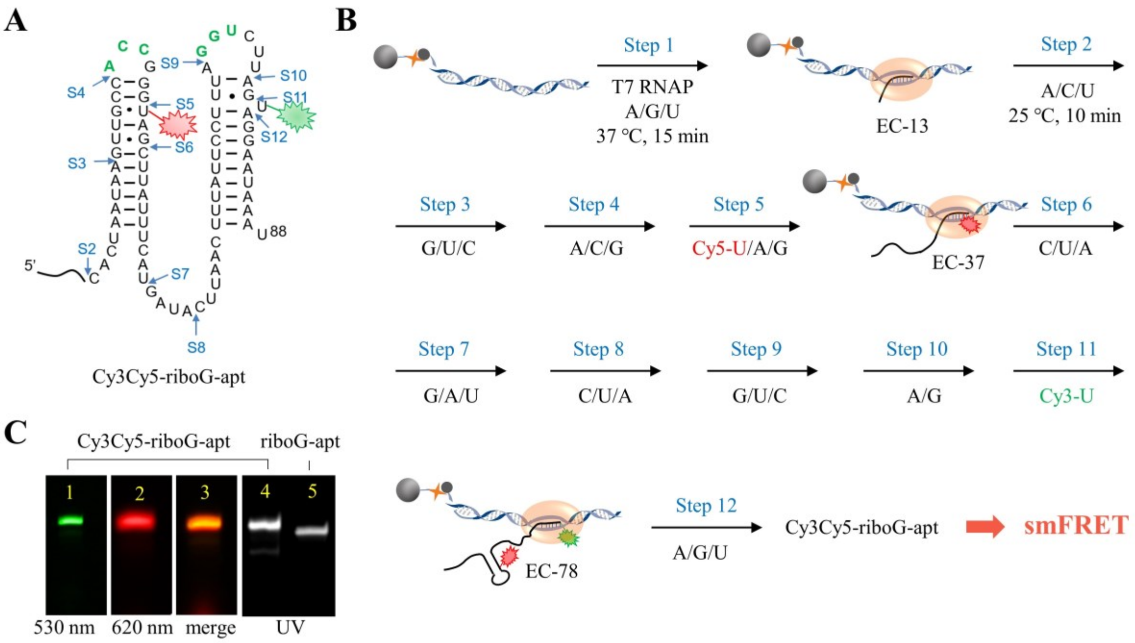


**Figure 2—figure supplement 1 | The schematic procedure of preparing Cy3Cy5-riboG-apt by 12 step-PLOR reaction for smFRET study.** A, The secondary structure of riboG-apt. The positions of donor (Cy3) and acceptor (Cy5) are shown by green and red sparkles at U78 and U35, respectively. The sites started at each step of the PLOR-synthesis were marked by blue arrows. B, The scheme of 12-step PLOR reaction for preparing Cy3Cy5-riboG-apt for smFRET study. The reagent usages for the 12-step reaction are listed in Supplementary Table S3. C, The PAGE images of Cy3Cy5-riboG-apt. The Lanes 1, 2 and 4 were irradiated by 530 nm, 620 nm fluorescence and 260 nm UV, respectively. Lane 3 is the merged image of lanes 1 and 2. The unlabeled counterpart, riboG-apt was loaded at lane 5. Cy3Cy5-riboG-apt (lane 4) migrated more slowly than riboG-apt (lane 5), indicating the bulky fluorophores were successfully introduced to Cy3Cy5-riboG-apt.

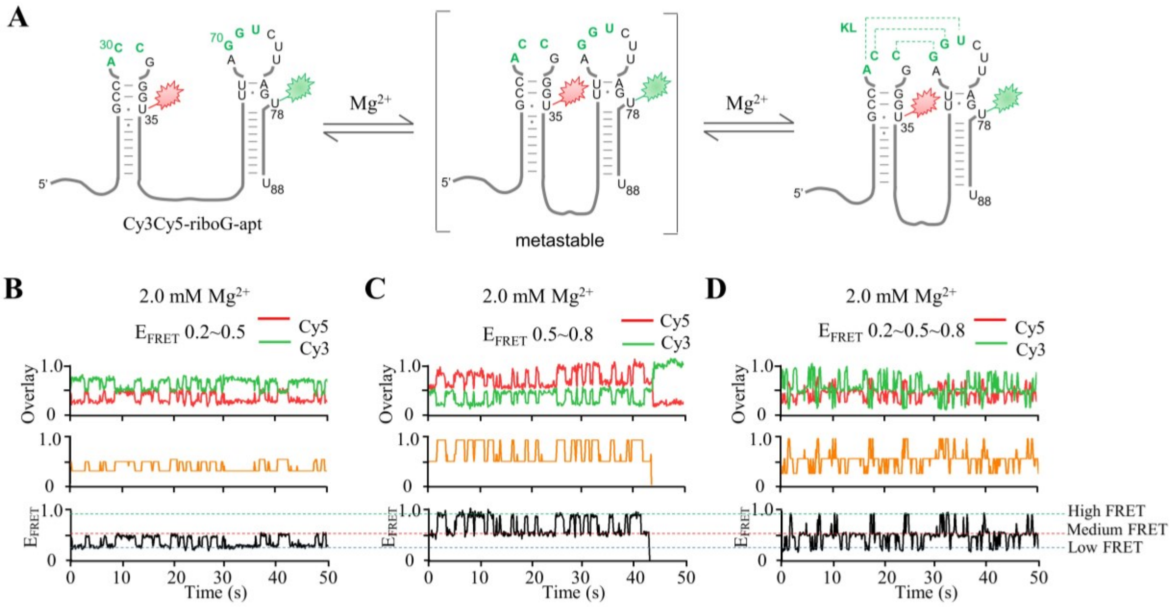


**Figure 2—figure supplement 2 │ smFRET measurements of Cy3Cy5-riboG-apt at 2.0 mM $Mg^{2+}$.** A, The secondary structures of the unfolded (left), pre-folded (medium) and folded (right) states of riboG-apt. B–D, HMM analysis of the representative single-molecule trajectories with the transitions between $E_{FRET}$ ~ 0.2 and $E_{FRET}$ ~ 0.5 (B), $E_{FRET}$ ~ 0.5 and ~ 0.8 (C), and among $E_{FRET}$ ~ 0.2, 0.5 and 0.8 (D). The time resolution for smFRET is 100 ms.

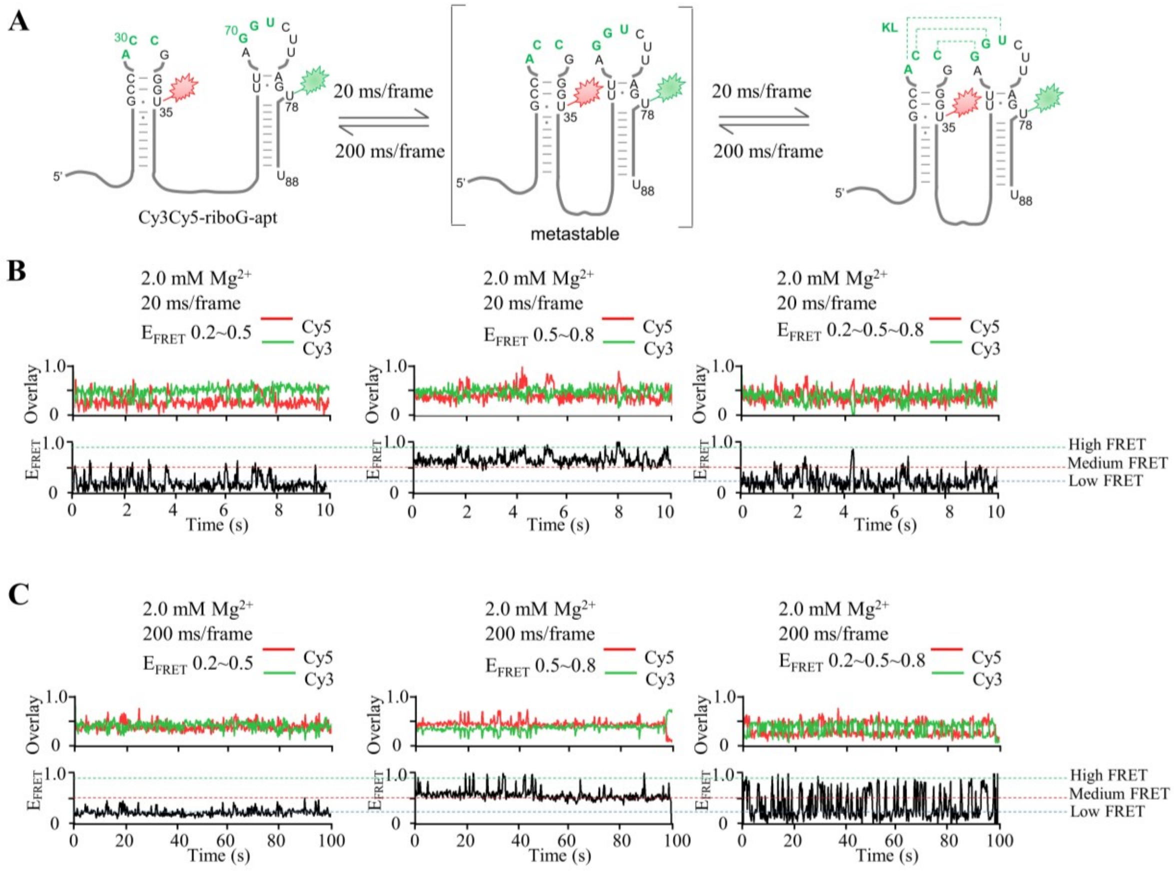


**Figure 2—figure supplement 3 │ smFRET measurements of Cy3Cy5-riboG-apt at different time resolution.** A, The secondary structures of the unfolded (left), pre-folded (medium) and folded (right) states of riboG-apt. B–C, The representative single-molecule trajectories and FRET curves with dynamic transitions. The data collection was performed with time resolution of 20 ms (B) and 200 ms (C).

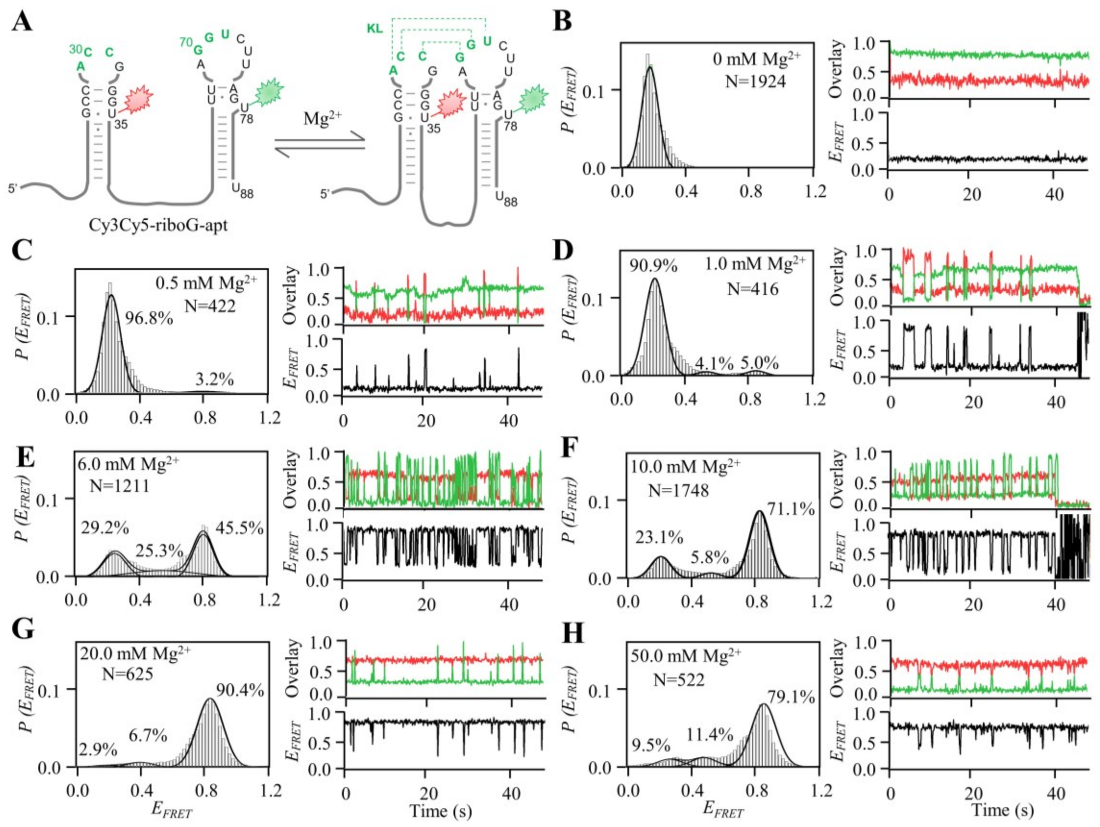


**Figure 2—figure supplement 4 │ smFRET measurements of Cy3Cy5-riboG-apt at 0–50.0 mM $Mg^{2+}$.** A, The secondary structures of the unfolded (left) and folded (right) states of riboG-apt. B–H, smFRET histograms, representative single-molecule trajectories and FRET curves of Cy3Cy5-riboG-apt at 0 mM $Mg^{2+}$ (B), 0.5 mM $Mg^{2+}$ (C), 1.0 mM $Mg^{2+}$ (D), 6.0 mM $Mg^{2+}$ (E), 10.0 mM $Mg^{2+}$ (F), 20.0 mM $Mg^{2+}$ (G), and 50.0 mM $Mg^{2+}$ (H).

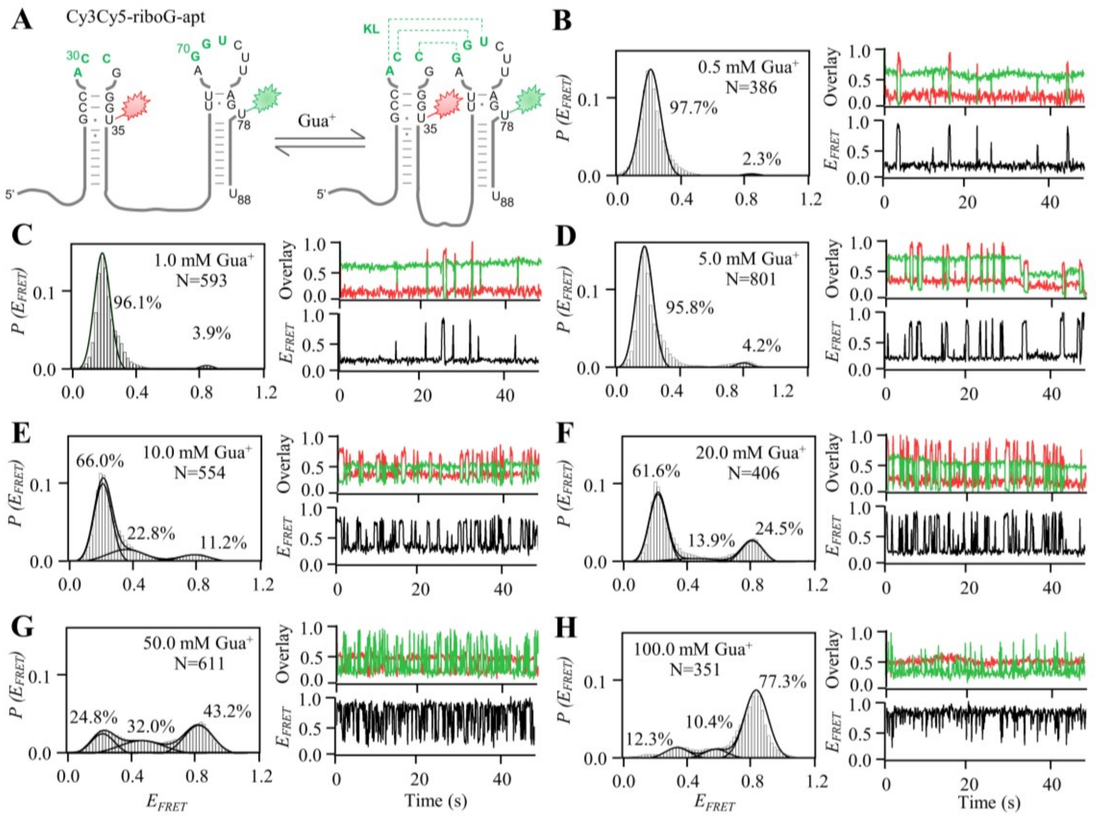

**Figure 2—figure supplement 5 │ smFRET measurements of Cy3Cy5-riboG-apt at 0.5 – 100.0 mM Gua$^+$.** A, The secondary structures of the unfolded (left) and folded (right) states of riboG-apt. B–H, smFRET histograms, representative single-molecule trajectories and FRET curves of Cy3Cy5-riboG-apt at 0.5 mM Gua$^+$ (B), 1.0 mM Gua$^+$ (C), 5.0 mM Gua$^+$ (D), 10.0 mM Gua$^+$ (E), 20.0 mM Gua$^+$ (F), 50.0 mM Gua$^+$ (G), and 100.0 mM Gua$^+$ (H).

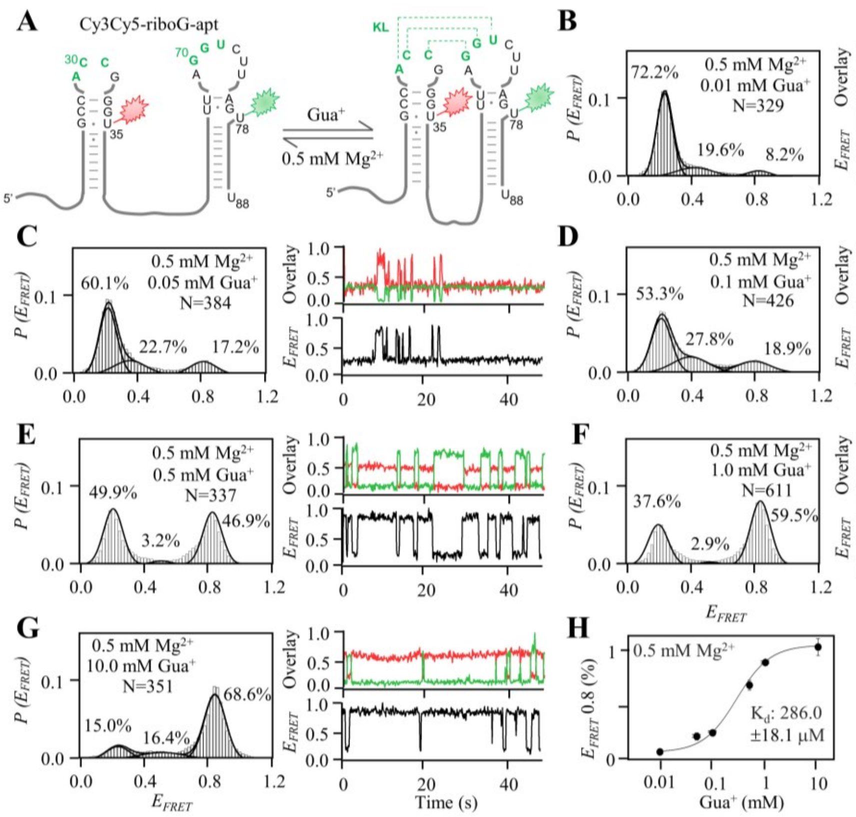


**Figure 2—figure supplement 6 │ smFRET measurements of Cy3Cy5-riboG-apt at 0.01–10.0 mM $Gua^+$ in the presence of 0.5 mM $Mg^{2+}$.** A, The secondary structures of the unfolded (left) and the folded (right) states of riboG-apt. B–G, smFRET histograms, representative single-molecule trajectories and FRET curves of Cy3Cy5-riboG-apt at 0.01 mM $Gua^+$ (B), 0.05 mM $Gua^+$ (C), 0.1 mM $Gua^+$ (D), 0.5 mM $Gua^+$ (E), 1.0 mM $Gua^+$ (F), and 10.0 mM $Gua^+$ (G) in the presence of 0.5 mM $Mg^{2+}$. H, The percentages of the folded state ($E_{FRET} \sim 0.8$) of Cy3Cy5-riboG-apt were plotted with the concentrations of $Gua^+$ at 0.5 mM $Mg^{2+}$, with an apparent $K_d$ of 286.0 ± 18.1 µM in three independent experiments.

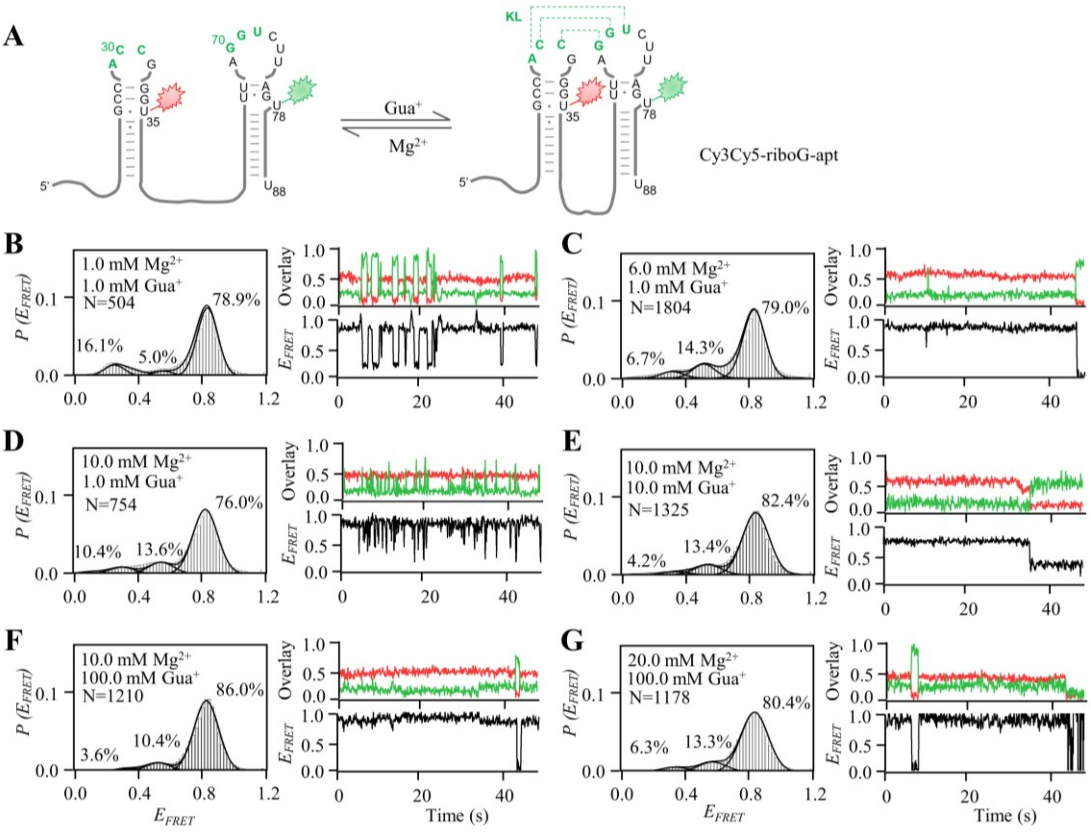


**Figure 2—figure supplement 7 │smFRET measurements of Cy3Cy5-riboG-apt at different Gua$^{+}$ and Mg$^{2+}$.** A, The secondary structures of the unfolded (left) and the folded (right) states of riboG-apt. B–G, smFRET histograms, representative single-molecule trajectories and FRET curves of Cy3Cy5-riboG-apt at 1.0 mM Gua$^{+}$ and 1.0 mM Mg$^{2+}$ (B), 1.0 mM Gua$^{+}$ and 6.0 mM Mg$^{2+}$ (C), 1.0 mM Gua$^{+}$ and 10.0 mM Mg$^{2+}$ (D), 10.0 mM Gua$^{+}$ and 10.0 mM Mg$^{2+}$ (E), 100.0 mM Gua$^{+}$ and 10.0 mM Mg$^{2+}$ (F), and 100.0 mM Gua$^{+}$ and 20.0 mM Mg$^{2+}$ (G).

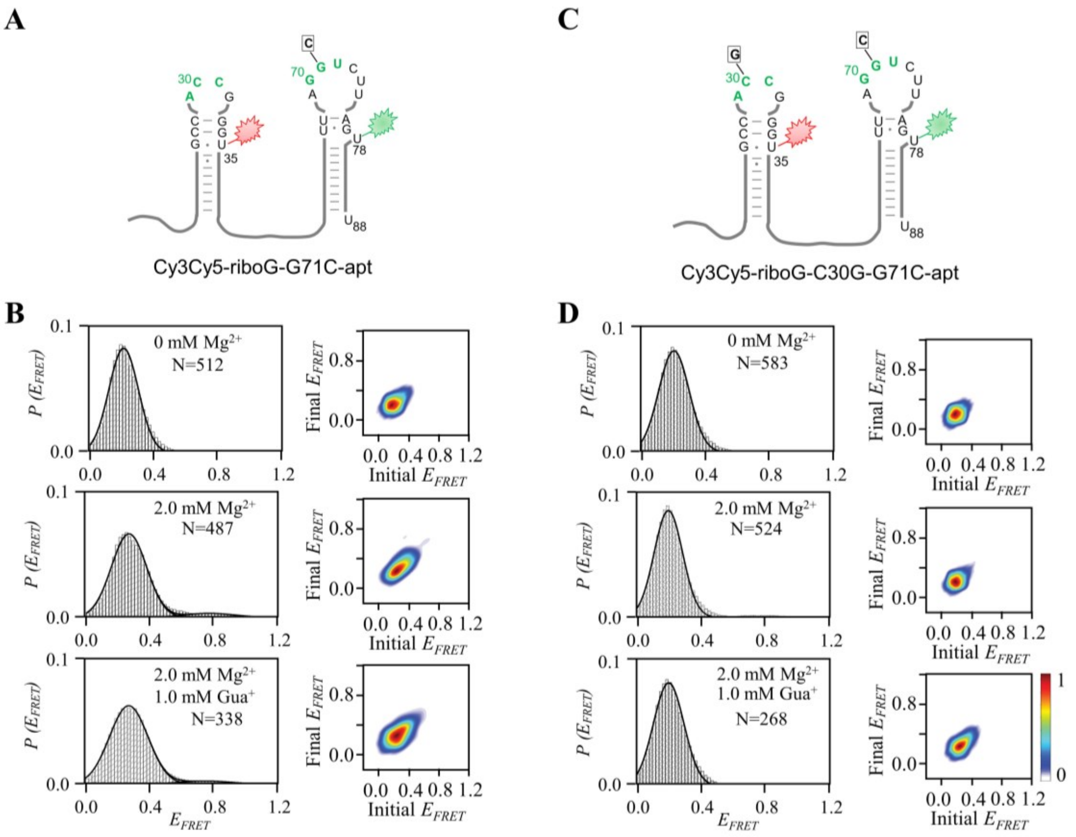


**Figure 2—figure supplement 8 │smFRET measurements of Cy3Cy5-riboG-G71C-apt and Cy3Cy5-riboG-C30G-G71C-apt.** A, The secondary structure of Cy3Cy5-riboG-G71C-apt. B, smFRET histograms and transition density plots for Cy3Cy5-riboG-G71C-apt at 0 mM $Mg^{2+}$, at 2.0 mM $Mg^{2+}$, and at 2.0 mM $Mg^{2+}$ and 1.0 mM $Gua^{+}$. C, The secondary structure of Cy3Cy5-riboG-C30G-G71C-apt. D, smFRET histograms and transition density plots for Cy3Cy5-riboG-C30G-G71C-apt at 0 mM $Mg^{2+}$, at 2.0 mM $Mg^{2+}$, and at 2.0 mM $Mg^{2+}$ and 1.0 mM $Gua^{+}$.

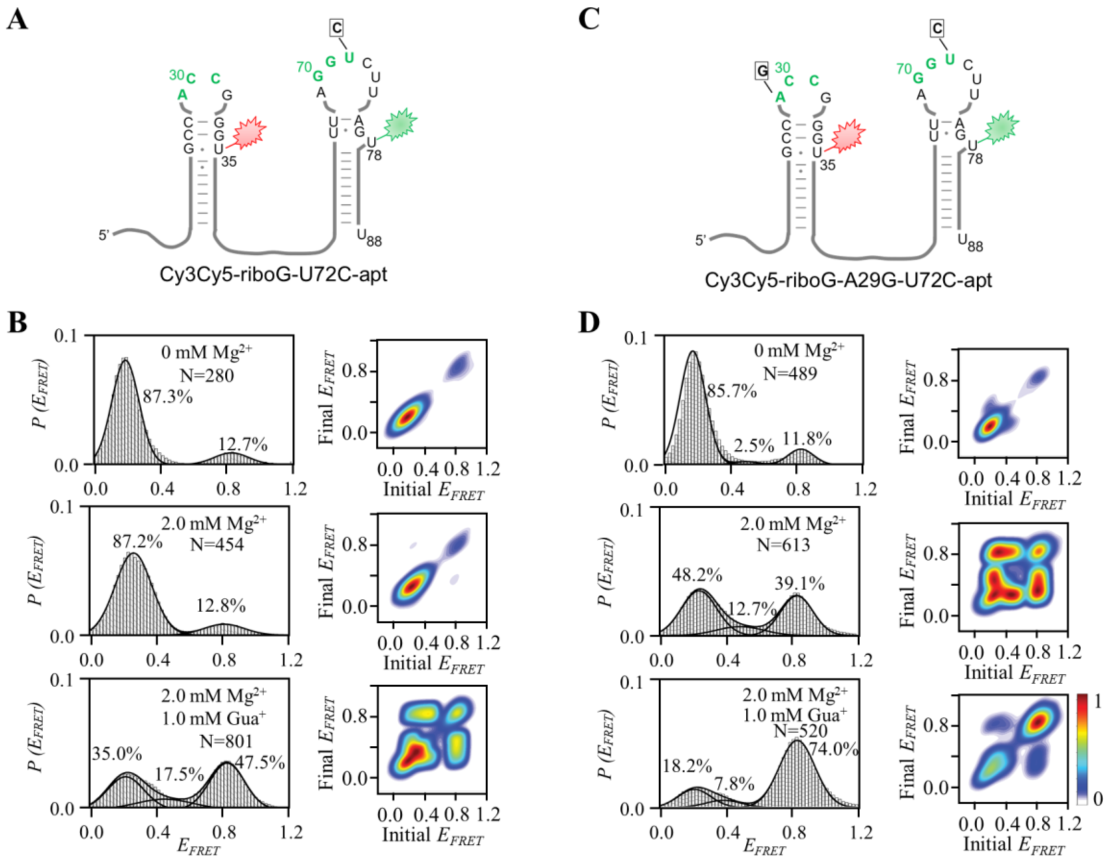


**Figure 2—figure supplement 9 ❘ smFRET measurements of Cy3Cy5-riboG-U72C-apt and Cy3Cy5-riboG-A29G-U72C-apt.** A, The secondary structure of Cy3Cy5-riboG-U72C-apt. B, smFRET histograms and transition density plots for Cy3Cy5-riboG-U72C-apt at 0 mM $Mg^{2+}$, at 2.0 mM $Mg^{2+}$, and at 2.0 mM $Mg^{2+}$ and 1.0 mM $Gua^{+}$. C, The secondary structure of Cy3Cy5-riboG-A29G-U72C-apt. D, smFRET histograms and transition density plots for Cy3Cy5-riboG-A29G-U72C-apt at 0 mM $Mg^{2+}$, at 2.0 mM $Mg^{2+}$, and at 2.0 mM $Mg^{2+}$ and 1.0 mM $Gua^{+}$.

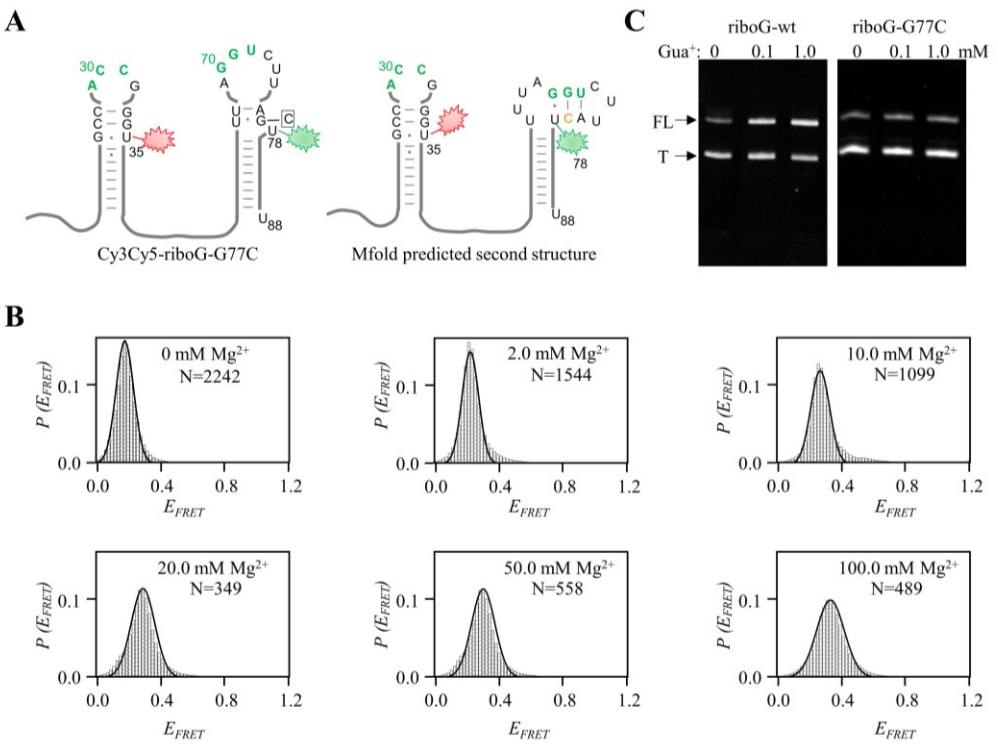


**Figure 2—figure supplement 10 │ The G77C mutation perturbs the folding and function of riboG-apt.** A, The secondary structures of Cy3Cy5-riboG-G77C-apt, with the structure predicted by Mfold on the right. B, smFRET histograms of Cy3Cy5-riboG-G77C-apt at 0–100.0 mM $Mg^{2+}$. C, Denaturing PAGE images of transcription termination assays of WT (left) and riboG-G77C, respectively. The FL and T represent the full-length and terminated RNA.

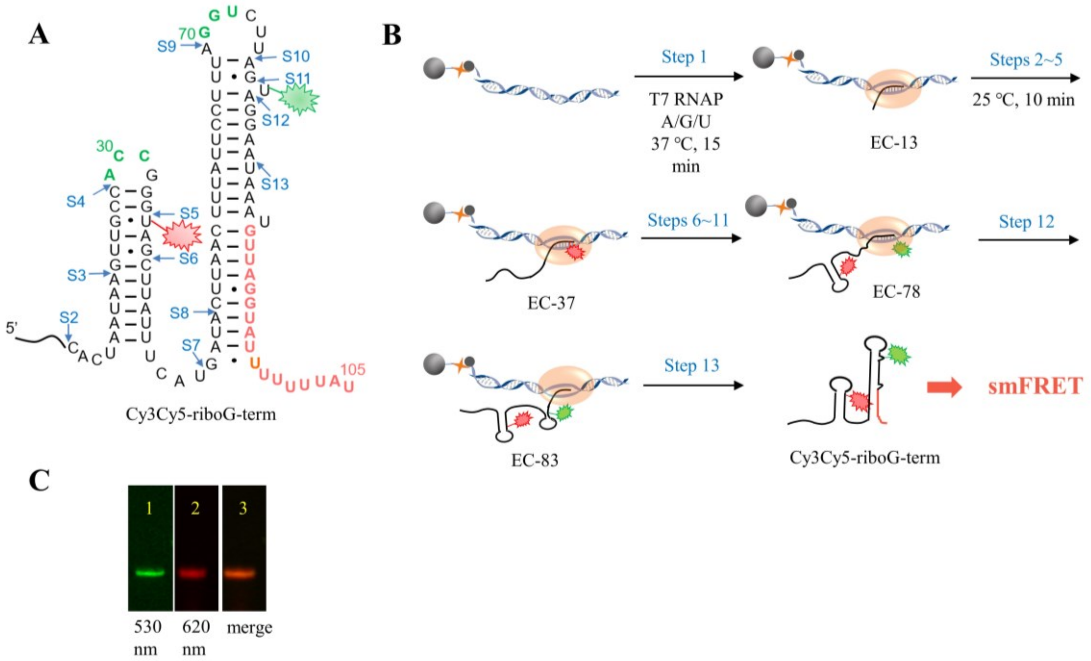


**Figure 3—figure supplement 1 | The schematic procedure of preparing Cy3Cy5-riboG-term by 13 step-PLOR reaction for smFRET study.** A, The secondary structure of riboG-term. The positions of donor (Cy3) and acceptor (Cy5) are shown by green and red sparkles at U78 and U35, respectively. The sites started at each step of the PLOR-synthesis were marked by blue arrows. B, 13-step PLOR reaction for preparing Cy3Cy5-riboG-term for smFRET study. The reagent usages for the 13-step reaction are listed in Supplementary Table S4. C, The PAGE images of Cy3Cy5-riboG-term. The Lanes 1 and 2 were irradiated by 530 nm and 620 nm fluorescence, respectively. Lane 3 is the merged image of lanes 1 and 2.

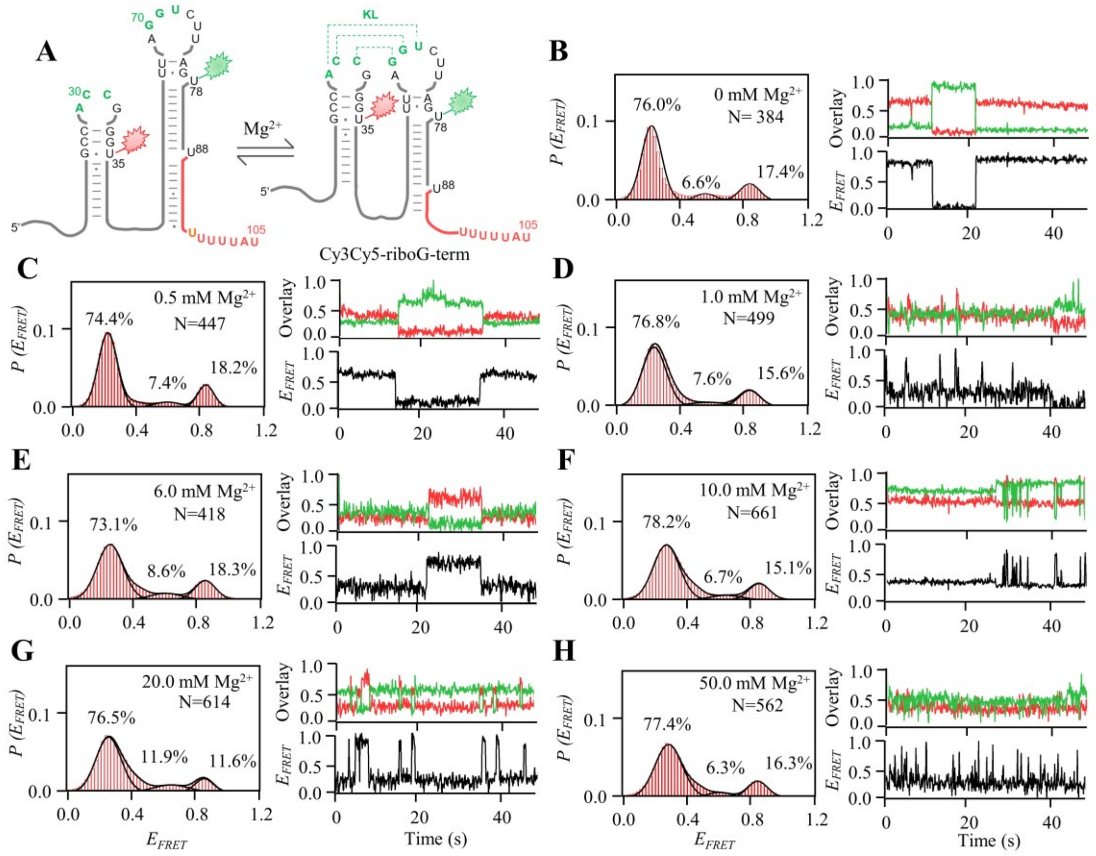


**Figure 3—figure supplement 2 │ smFRET measurements of Cy3Cy5-riboG-term at 0–50.0 mM Mg$^{2+}$.** A, The secondary structures of the unfolded (left) and folded (right) states of riboG-term. B–H, smFRET histograms, dynamic single-molecule trajectories and FRET curves of Cy3Cy5-riboG-term at 0 mM Mg$^{2+}$ (B), 0.5 mM Mg$^{2+}$ (C), 1.0 mM Mg$^{2+}$ (D), 6.0 mM Mg$^{2+}$ (E), 10.0 mM Mg$^{2+}$ (F), 20.0 mM Mg$^{2+}$ (G), and 50.0 mM Mg$^{2+}$ (H). Approximately 1.4%, 1.1%, 6.2%, 7.4%, 10.3%, 6.2%, and 2.4% of dynamic single-molecule trajectories were detected in (B), (C), (D), (E), (F), (G), and (H), respectively.

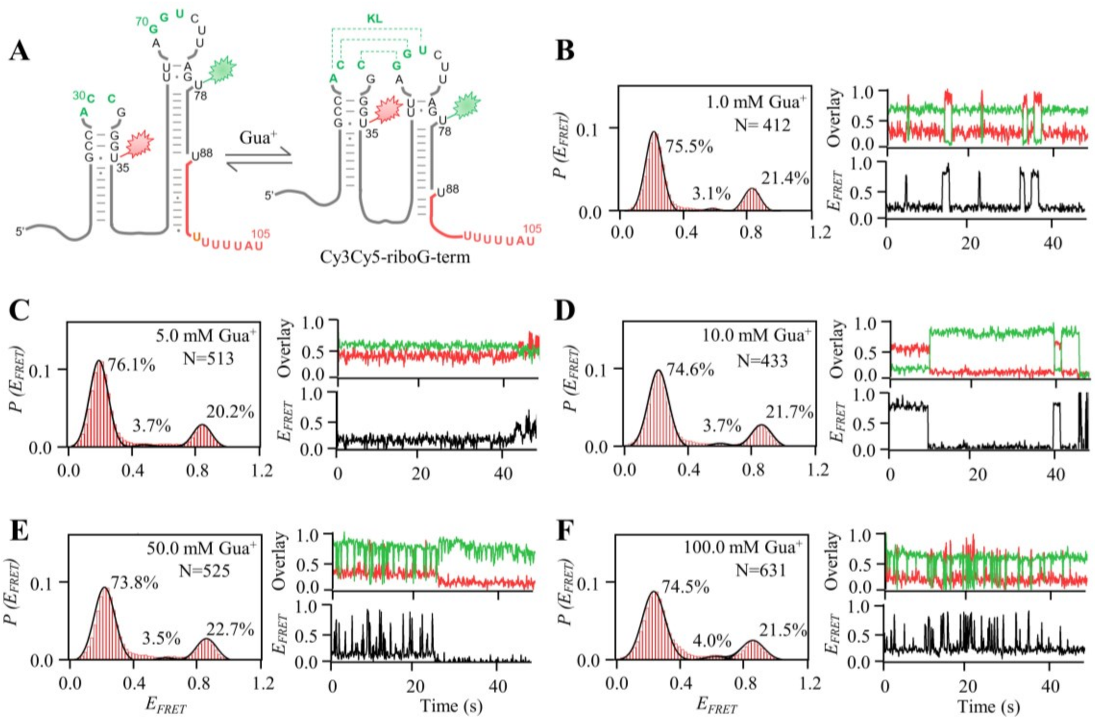


**Figure 3—figure supplement 3 │smFRET measurements of Cy3Cy5-riboG-term at 1.0 –100.0 mM Gua⁺.** A, The secondary structures of the unfolded (left) and folded (right) states of riboG-term. B–F, smFRET histograms, dynamic single-molecule trajectories and FRET curves of Cy3Cy5-riboG-term at 1.0 mM Gua⁺ (B), 5.0 mM Gua⁺ (C), 10.0 mM Gua⁺ (D), 50.0 mM Gua⁺ (E) and 100.0 mM Gua⁺ (F). Approximately 1.3%, 3.4%, 3.1%, 3.0%, and 11.5% of dynamic single-molecule trajectories were detected in (B), (C), (D), (E), and (F), respectively.

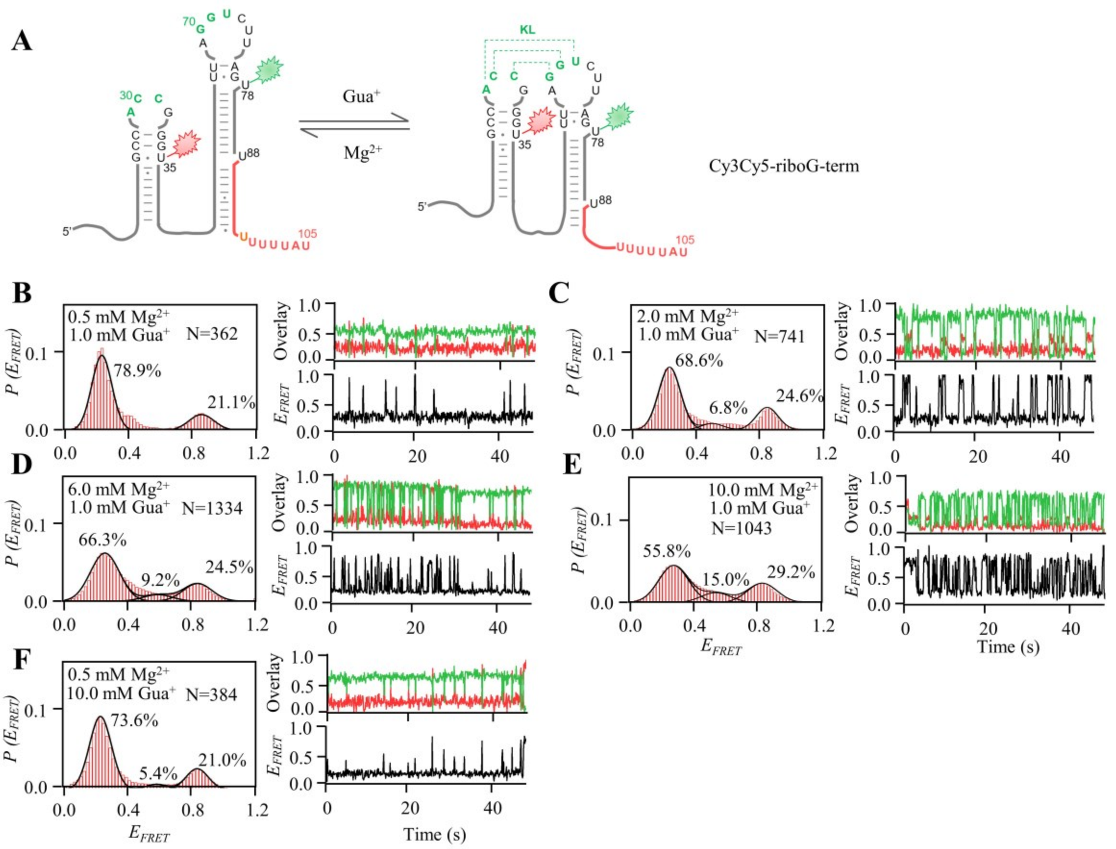


**Figure 3—figure supplement 4 │smFRET measurements of Cy3Cy5-riboG-term at different Gua$^{+}$ and Mg$^{2+}$.** A, The secondary structures of the unfolded (left) and folded (right) states of riboG-term. B–F, smFRET histograms, representative single-molecule trajectories and FRET curves of Cy3Cy5-riboG-term at 1.0 mM Gua$^{+}$ and 0.5 mM Mg$^{2+}$ (B), 1.0 mM Gua$^{+}$ and 2.0 mM Mg$^{2+}$ (C), 1.0 mM Gua$^{+}$ and 6.0 mM Mg$^{2+}$ (D), 1.0 mM Gua$^{+}$ and 10.0 mM Mg$^{2+}$ (E), and 10.0 mM Gua$^{+}$ and 0.5 mM Mg$^{2+}$ (F).

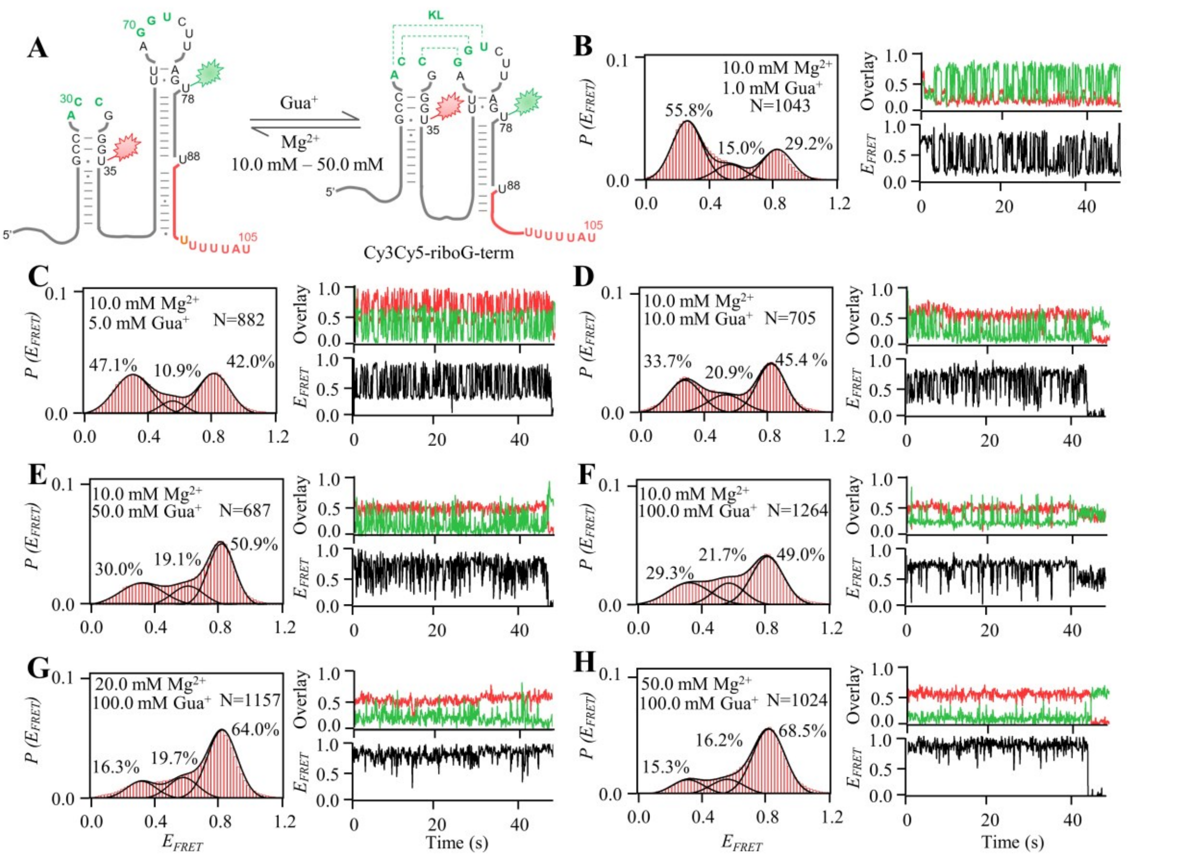


**Figure 3—figure supplement 5 | smFRET measurements of Cy3Cy5-riboG-term at different Gua$^{+}$ and Mg$^{2+}$.** A, The secondary structures of the unfolded (left) and folded (right) states of riboG-term. B–H, smFRET histograms, representative single-molecule trajectories and FRET curves of Cy3Cy5-riboG-term at 1.0 mM Gua$^{+}$ and 10.0 mM Mg$^{2+}$ (B), 5.0 mM Gua$^{+}$ and 10.0 mM Mg$^{2+}$ (C), 10.0 mM Gua$^{+}$ and 10.0 mM Mg$^{2+}$ (D), 50.0 mM Gua$^{+}$ and 10.0 mM Mg$^{2+}$ (E), 100.0 mM Gua$^{+}$ and 10.0 mM Mg$^{2+}$ (F), 100.0 mM Gua$^{+}$ and 20.0 mM Mg$^{2+}$ (G), and 100.0 mM Gua$^{+}$ and 50.0 mM Mg$^{2+}$ (H).

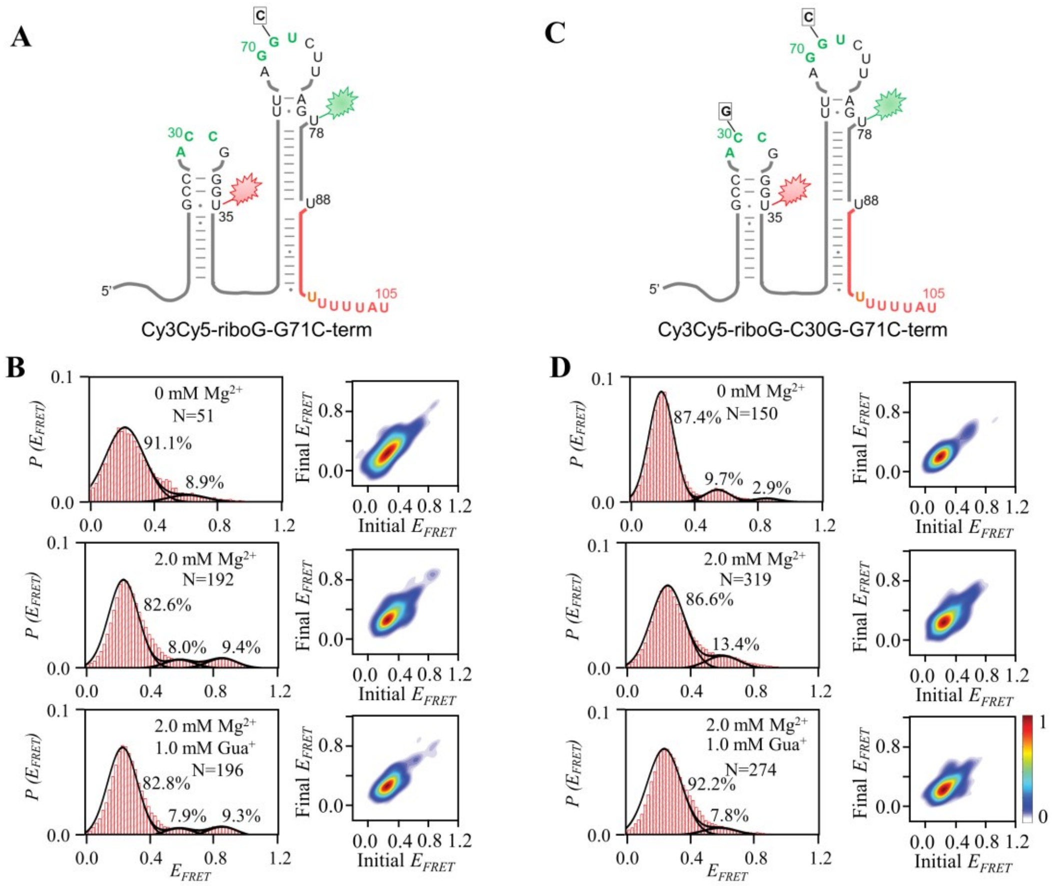


**Figure 3—figure supplement 6 │smFRET measurements for Cy3Cy5-riboG-G71C-term and Cy3Cy5-riboG-C30G-G71C-term.** A, The secondary structure of Cy3Cy5-riboG-G71C-term. B, smFRET histograms and transition density plots for Cy3Cy5-riboG-G71C-term at 0 mM $Mg^{2+}$, at 2.0 mM $Mg^{2+}$, and at 2.0 mM $Mg^{2+}$ and 1.0 mM $Gua^{+}$. C, The secondary structure of Cy3Cy5-riboG-C30G-G71C-term. D, smFRET histograms and transition density plots for Cy3Cy5-riboG-C30G-G71C-term at 0 mM $Mg^{2+}$, at 2.0 mM $Mg^{2+}$, and at 2.0 mM $Mg^{2+}$ and 1.0 mM $Gua^{+}$.

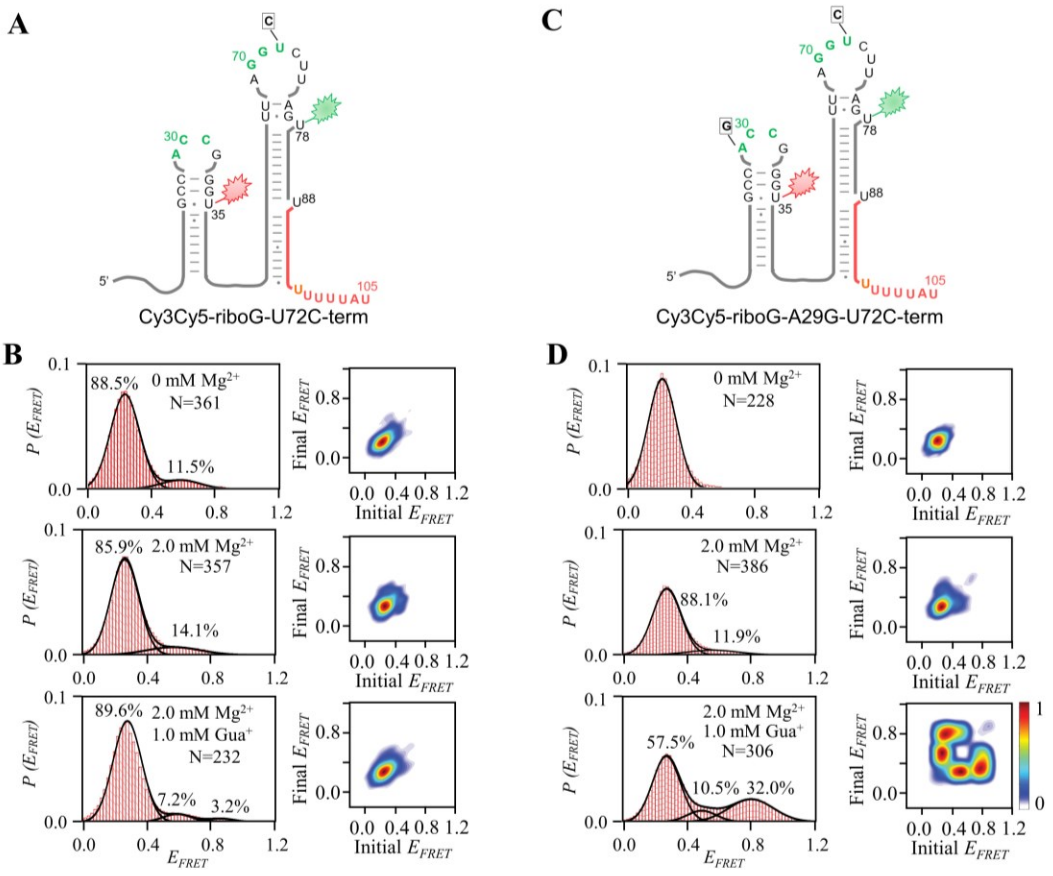


**Figure 3—figure supplement 7 | smFRET measurements for Cy3Cy5-riboG-U72C-term and Cy3Cy5-riboG-A29G-U72C-term.** A, The secondary structure of Cy3Cy5-riboG-U72C-term. B, smFRET histograms and transition density plots for Cy3Cy5-riboG-U72C-term at 0 mM $Mg^{2+}$, at 2.0 mM $Mg^{2+}$, and at 2.0 mM $Mg^{2+}$ and 1.0 mM $Gua^{+}$. C, The secondary structure of Cy3Cy5-riboG-A29G-U72C-term. D, smFRET histograms and transition density plots for Cy3Cy5-riboG-A29G-U72C-term at 0 mM $Mg^{2+}$, at 2.0 mM $Mg^{2+}$, and at 2.0 mM $Mg^{2+}$ and 1.0 mM $Gua^{+}$.

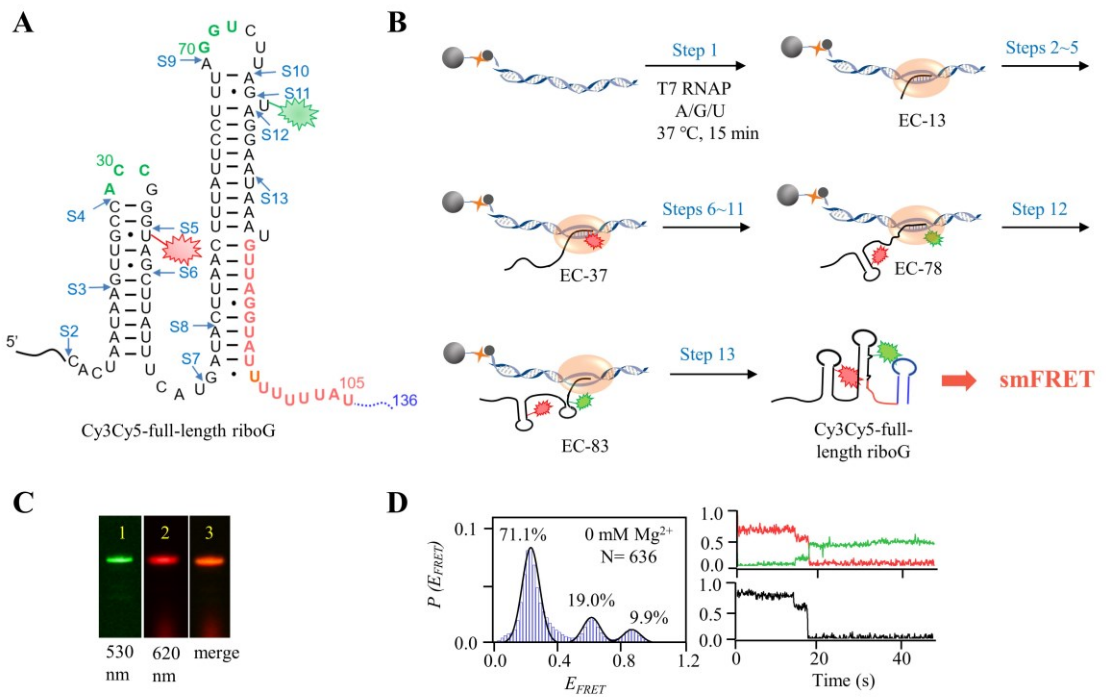


**Figure 4—figure supplement 1 | The schematic procedure of preparing Cy3Cy5-full-length riboG by 13 step-PLOR reaction for smFRET study.** A, The secondary structure of full-length riboG. The positions of donor (Cy3) and acceptor (Cy5) are shown by green and red sparkles at U78 and U35, respectively. The sites started at each step of the PLOR-synthesis were marked by blue arrows. B, 13-step PLOR reaction for preparing Cy3Cy5-full-length riboG for smFRET study. The reagent usages for the 13-step reaction are listed in Supplementary Table S5. C, The PAGE images of Cy3Cy5-full-length riboG. The Lanes 1 and 2 were irradiated by 530 nm and 620 nm fluorescence, respectively. Lane 3 is the merged image of lanes 1 and 2. D, smFRET study of Cy3Cy5-full-length riboG in the absence of $Mg^{2+}$. The smFRET histogram, dynamic single-molecule trajectory and FRET curve for Cy3Cy5-full-length riboG at 0 mM $Mg^{2+}$. Approximately 1.4% of dynamic single-molecule trajectories was detected.

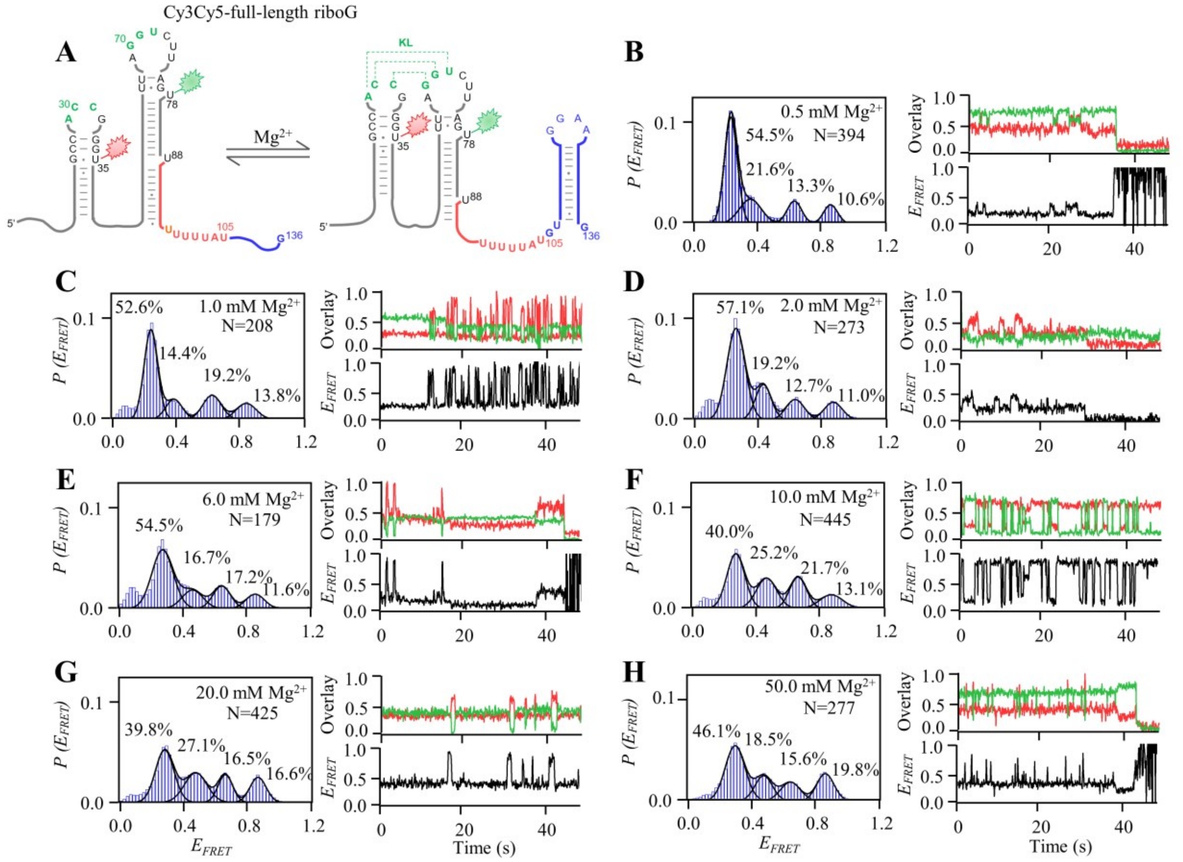

**Figure 4—figure supplement 2 │smFRET measurements of Cy3Cy5-full-length riboG at 0.5–50.0 mM $Mg^{2+}$.** A, The secondary structures of the unfolded (left) and folded (right) states of full-length riboG. B–H, smFRET histograms, dynamic single-molecule trajectories and FRET curves of Cy3Cy5-full-length riboG at 0.5 mM $Mg^{2+}$ (B), 1.0 mM $Mg^{2+}$ (C), 2.0 mM $Mg^{2+}$ (D), 6.0 mM $Mg^{2+}$ (E), 10.0 mM $Mg^{2+}$ (F), 20.0 mM $Mg^{2+}$ (G), and 50.0 mM $Mg^{2+}$ (H). Approximately 2.6%, 3.8%, 5.5%, 11.7%, 11.5%, 16.0%, and 28.9% of dynamic single-molecule trajectories were detected (B), (C), (D), (E), (F), (G), and (H), respectively.

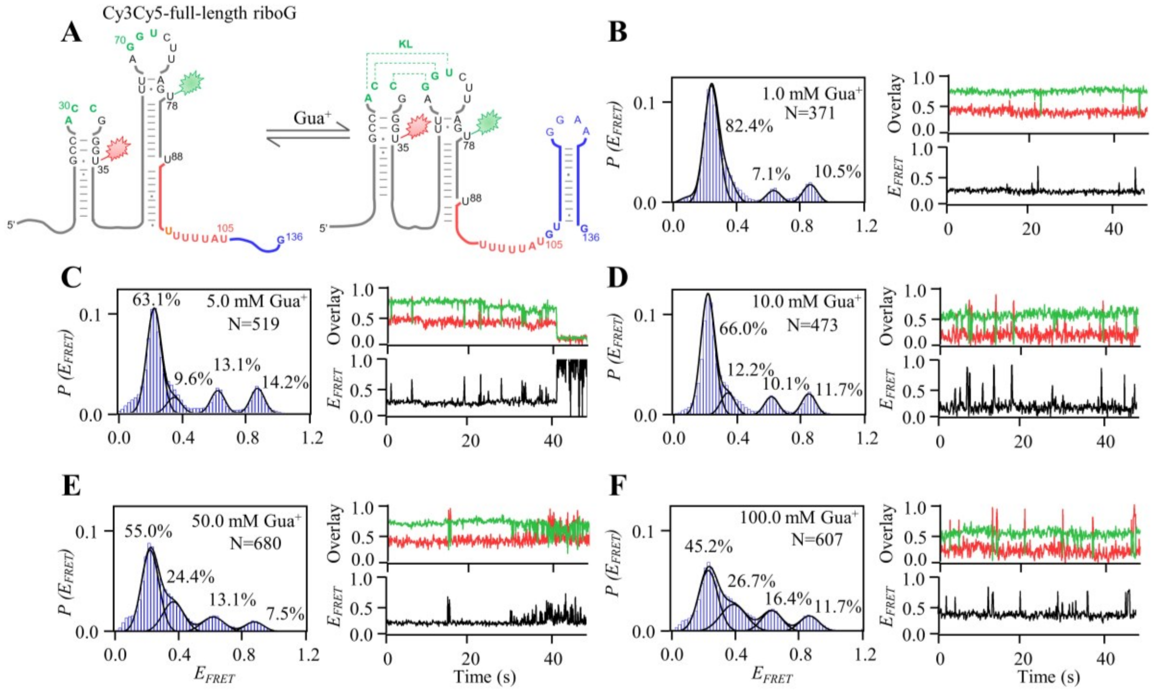


**Figure 4—figure supplement 3 │smFRET measurements of Cy3Cy5-full-length riboG at 1.0–100.0 mM Gua$^+$.** A, The secondary structures of the unfolded (left) and folded (right) states of full-length riboG. B–F, smFRET histograms, dynamic single-molecule trajectories and FRET curves of Cy3Cy5-full-length riboG at 1.0 mM Gua$^+$ (B), 5.0 mM Gua$^+$ (C), 10.0 mM Gua$^+$ (D), 50.0 mM Gua$^+$ (E) and 100.0 mM Gua$^+$ (F). Approximately 3.0%, 5.6%, 15.9%, 27.9%, and 46.6% of dynamic single-molecule trajectories were detected (B), (C), (D), (E), and (F), respectively.

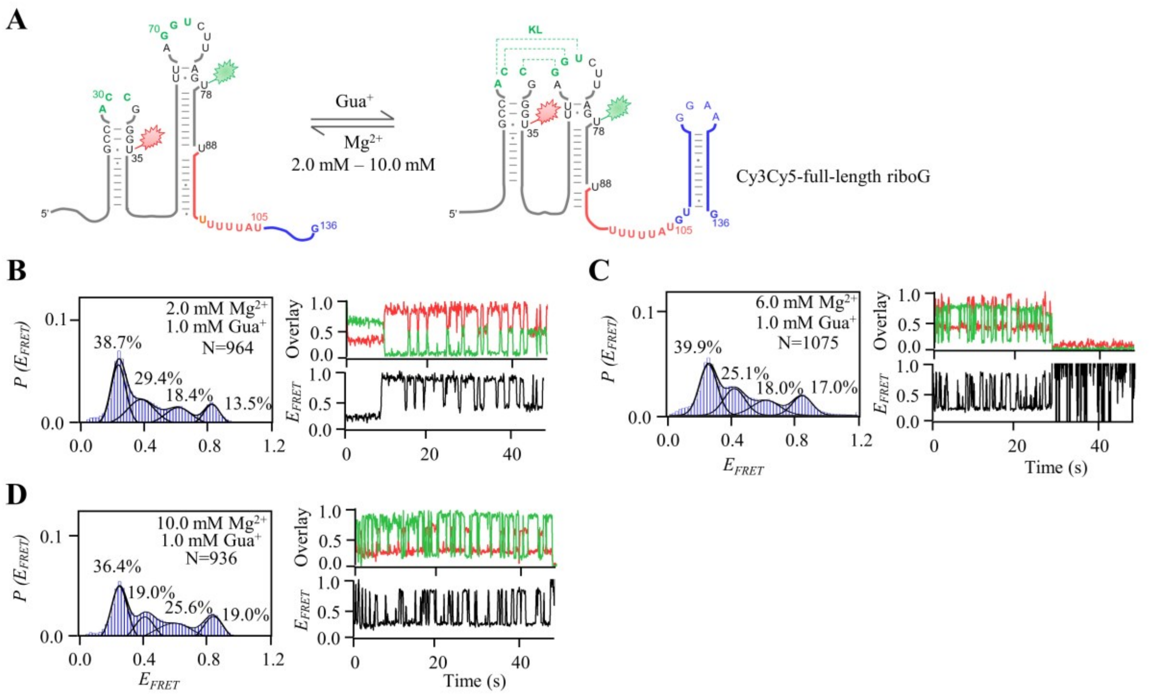


**Figure 4—figure supplement 4 | smFRET measurements of Cy3Cy5-full-length riboG at 1.0 mM Gua$^{+}$ and 2.0–10.0 mM Mg$^{2+}$.** A, The secondary structures of the unfolded (left) and folded (right) states of full-length riboG. B–D, smFRET histograms, dynamic single-molecule trajectories and FRET curves of Cy3Cy5-full-length riboG at 1.0 mM Gua$^{+}$ and 2 mM Mg$^{2+}$ (B), 1.0 mM Gua$^{+}$ and 6.0 mM Mg$^{2+}$ (C), and 1.0 mM Gua$^{+}$ and 10.0 mM Mg$^{2+}$ (D). Approximately 44.3%, 59.5%, and 66.5% of dynamic single-molecule trajectories were detected (B), (C), and (D), respectively.

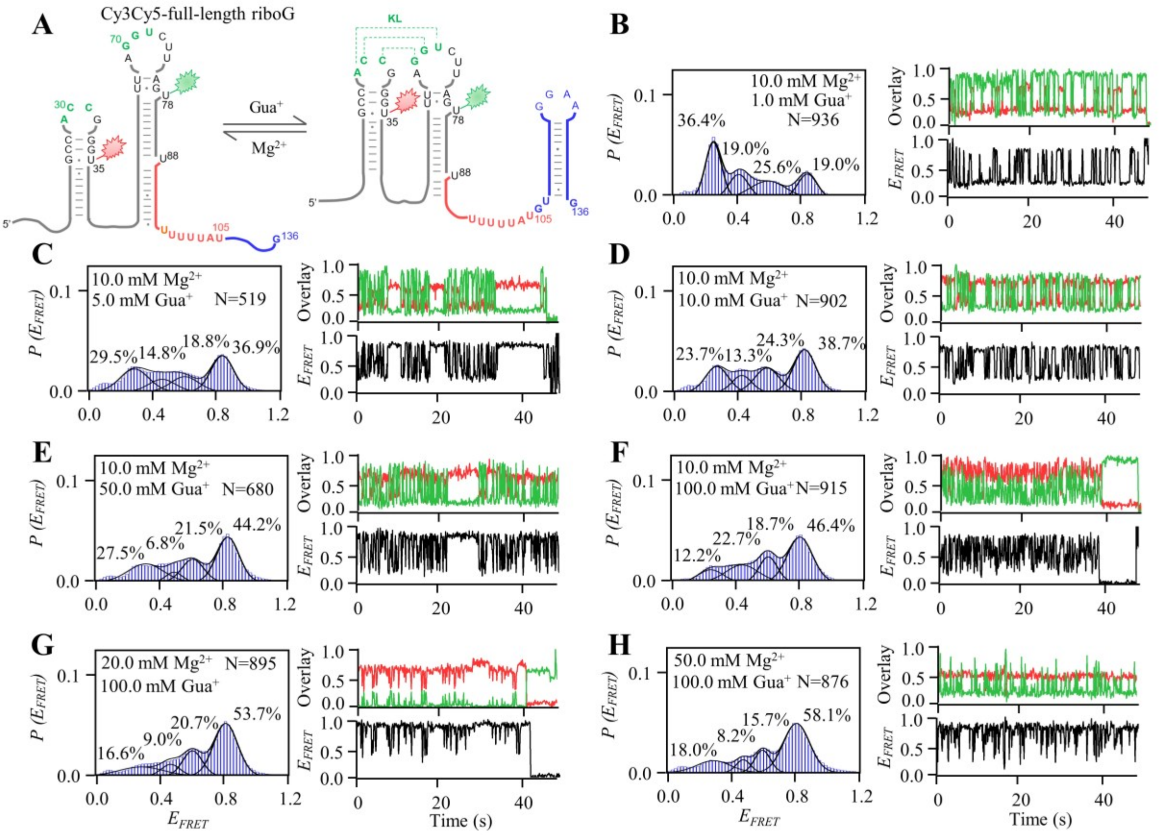


**Figure 4—figure supplement 5 │smFRET measurements of Cy3Cy5-full-length riboG at different Gua$^{+}$ and Mg$^{2+}$.** A, The secondary structures of the unfolded (left) and folded (right) states of full-length riboG. B–H, smFRET histograms, dynamic single-molecule trajectories and FRET curves of Cy3Cy5-full-length riboG at 1.0 mM Gua$^{+}$ and 10.0 mM Mg$^{2+}$ (B), 5.0 mM Gua$^{+}$ and 10.0 mM Mg$^{2+}$ (C), 10.0 mM Gua$^{+}$ and 10.0 mM Mg$^{2+}$ (D), 50.0 mM Gua$^{+}$ and 10.0 mM Mg$^{2+}$ (E), 100.0 mM Gua$^{+}$ and 10.0 mM Mg$^{2+}$ (F), 100.0 mM Gua$^{+}$ and 20.0 mM Mg$^{2+}$ (G), and 100.0 mM Gua$^{+}$ and 50.0 mM Mg$^{2+}$ (H). Approximately 66.5%, 83.4%, 65.7%, 79.6%, 66.9%, 63.9%, and 59.9% of dynamic single-molecule trajectories were detected (B), (C), (D), (E), (F), (G), and (H), respectively.

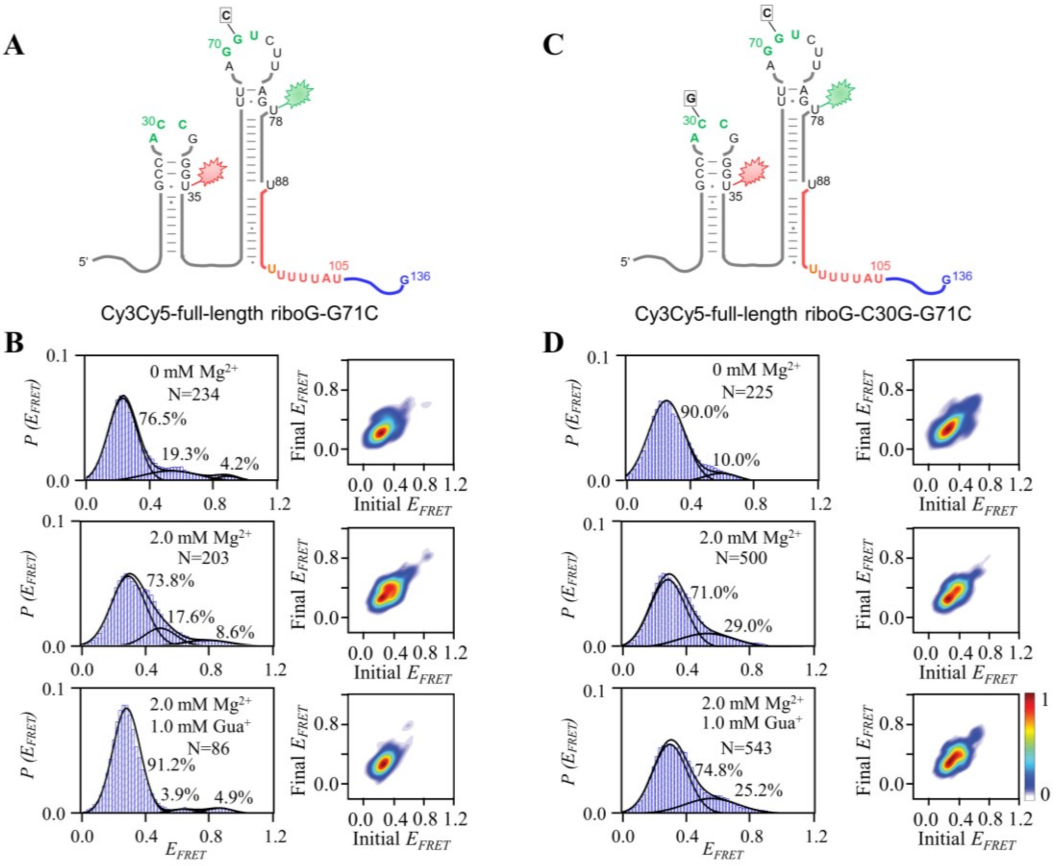


**Figure 4—figure supplement 6 │smFRET measurements for Cy3Cy5-full-length riboG-G71C and Cy3Cy5-full-length riboG-C30G-G71C.** A, The secondary structure of Cy3Cy5-full-length riboG-G71C. B, smFRET histograms and transition density plots for Cy3Cy5-full-length riboG-G71C at 0 mM $Mg^{2+}$, at 2.0 mM $Mg^{2+}$, and at 2.0 mM $Mg^{2+}$ and 1.0 mM $Gua^{+}$. C, The secondary structure of Cy3Cy5-full-length riboG-C30G-G71C. D, smFRET histograms and transition density plots for Cy3Cy5-full-length riboG-C30G-G71C at 0 mM $Mg^{2+}$, at 2.0 mM $Mg^{2+}$, and at 2.0 mM $Mg^{2+}$ and 1.0 mM $Gua^{+}$.

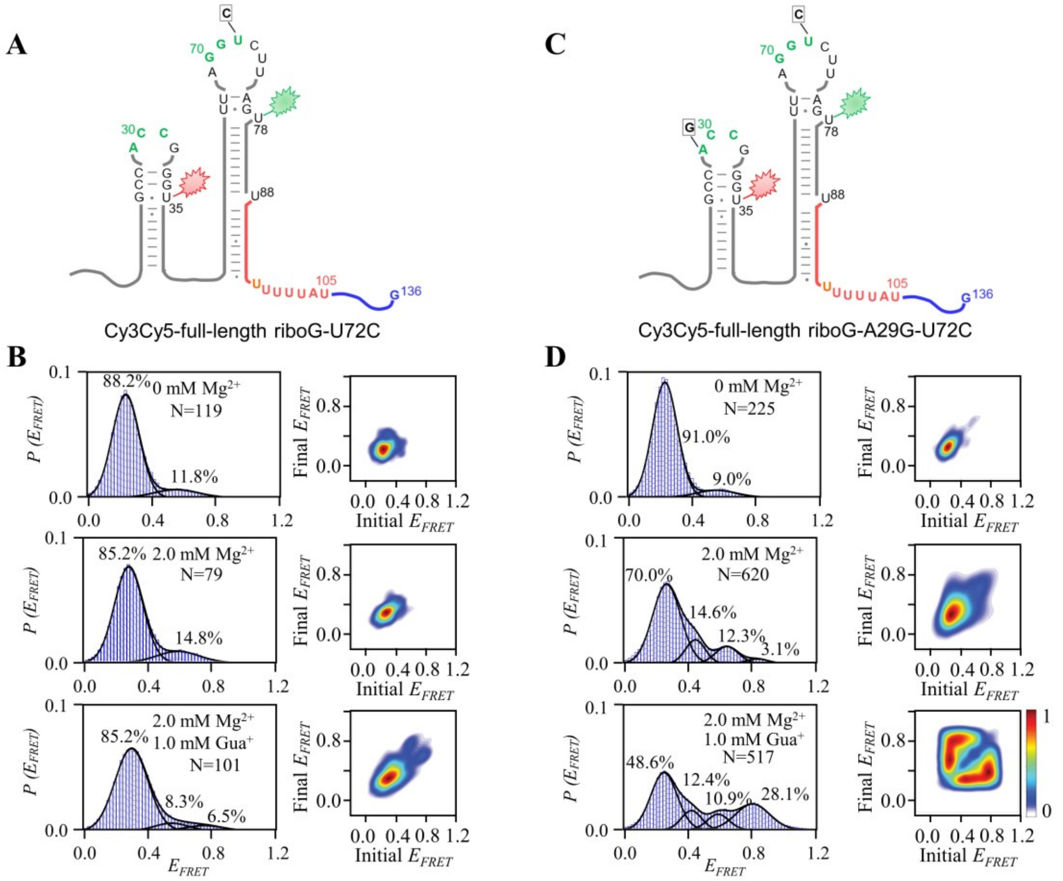


**Figure 4—figure supplement 7 | smFRET measurements for Cy3Cy5-full-length riboG-U72C and Cy3Cy5-full-length riboG-A29G-U72C.** A, The secondary structure of Cy3Cy5-full-length riboG-U72C. B, smFRET histograms and transition density plots for Cy3Cy5-full-length riboG-U72C at 0 mM $Mg^{2+}$, at 2.0 mM $Mg^{2+}$, and at 2.0 mM $Mg^{2+}$ and 1.0 mM $Gua^{+}$. C, The secondary structure of Cy3Cy5-full-length riboG-A29G-U72C. D, smFRET histograms and transition density plots for Cy3Cy5-full-length riboG-A29G-U72C at 0 mM $Mg^{2+}$, at 2.0 mM $Mg^{2+}$, and at 2.0 mM $Mg^{2+}$ and 1.0 mM $Gua^{+}$.

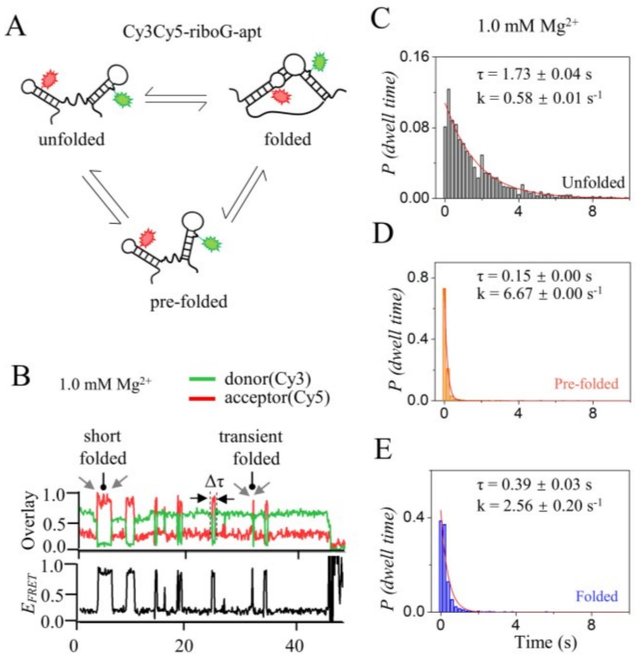


**Figure 5—figure supplement 1 │ Kinetics analysis of the unfolded, pre-folded and folded states of riboG-apt at 1.0 mM $Mg^{2+}$.** A, The structural switching of the unfolded, pre-folded and folded states of riboG-apt. The green and red sparkles represent Cy3 and Cy5, respectively. B, The representative single-molecule trajectories of riboG-apt at 1.0 mM $Mg^{2+}$. Δτ is the dwell time. C–E, Dwell time histograms for the unfolded (C), pre-folded (D) and folded (E) used for determination of transition rate constants at 1.0 mM $Mg^{2+}$. Plots were fit with exponential decay curves to extract conformation switch kinetics.

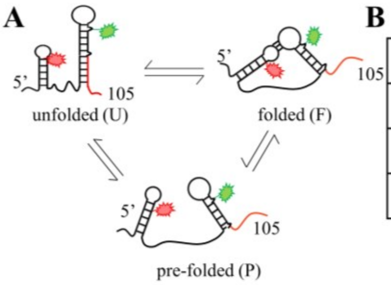


**B**

2.0 mM $Mg^{2+}$

| allosteric | U→P | P→U | U→F | F→U | P→F | F→P |
|---|---|---|---|---|---|---|
| n | 4 | 7 | 3 | 5 | 3 | 1 |
| t (s) | 3830 | | | | | |
| n/t (*$10^{-3}$ * $s^{-1}$) | 1.04 | 1.83 | 0.78 | 1.31 | 0.78 | 0.26 |

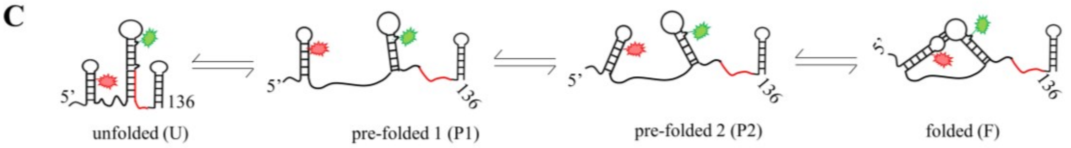


**D**

2.0 mM $Mg^{2+}$

| allosteric | U→P2 | U→P1 | U→F | P2→U | P2→P1 | P2→F | P1→U | P1→P2 | P1→F | F→U | F→P2 | F→P1 |
|---|---|---|---|---|---|---|---|---|---|---|---|---|
| n | 33 | 3 | 1 | 36 | 1 | 0 | 0 | 0 | 1 | 1 | 0 | 0 |
| t (s) | 6390 | | | | | | | | | | | |
| n/t (*$10^{-3}$ * $s^{-1}$) | 5.16 | 0.47 | 0.16 | 5.63 | 0.16 | 0.00 | 0.00 | 0.00 | 0.16 | 0.16 | 0.00 | 0.00 |

**Figure 5—figure supplement 2 | Kinetics analysis of isolated riboG-term and full-length riboG at 2.0 mM $Mg^{2+}$.** A, The structural switching of the unfolded (U), pre-folded (P) and folded states (F) of riboG-term. B, The transitions between two states of riboG-term at 2.0 mM $Mg^{2+}$. n: number of observed transition events; t: total observation time; n/t: frequency of transition between two states, and n/t is used to the rate constant of transitions (k). C, The structural switching of unfolded (U), pre-folded 1 (P1), pre-folded 2 (P2) and folded (F) states of full-length riboG. D, The transitions between two states of full-length riboG at the presence of 2.0 mM $Mg^{2+}$.

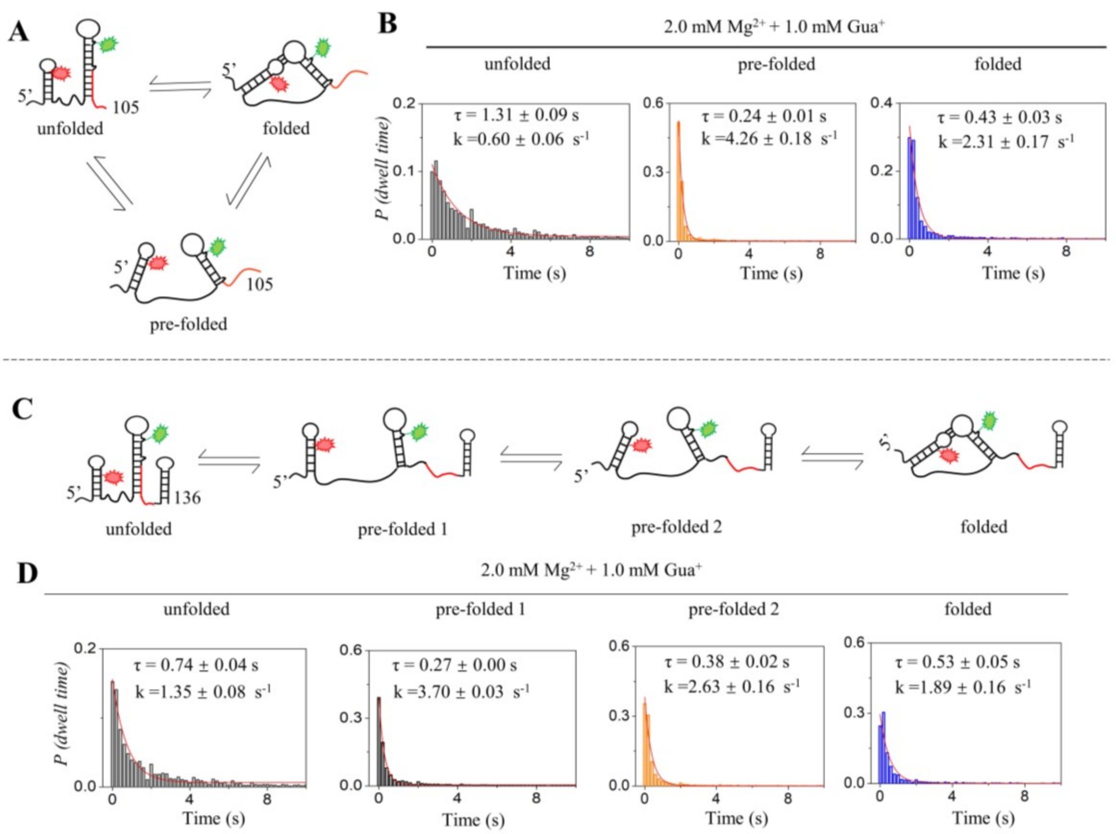


**Figure 5—figure supplement 3 │ Kinetics analysis of isolated riboG-term and full-length riboG at 2.0 mM $Mg^{2+}$ and 1.0 mM $Gua^{+}$.** A, The structural switching of the unfolded (U), pre-folded (P) and folded states (F) of riboG-term. B, The histograms of dwell time for the unfolded (gray), pre-folded (orange) and folded (blue) states of riboG-term. C, The structural switching of unfolded (U), pre-folded 1 (P1), pre-folded 2 (P2) and folded (F) states of full-length riboG. D, The histograms of dwell time for the unfolded (gray), pre-folded 1 (black), pre-folded 2 (orange) and folded (blue) states of full-length riboG.

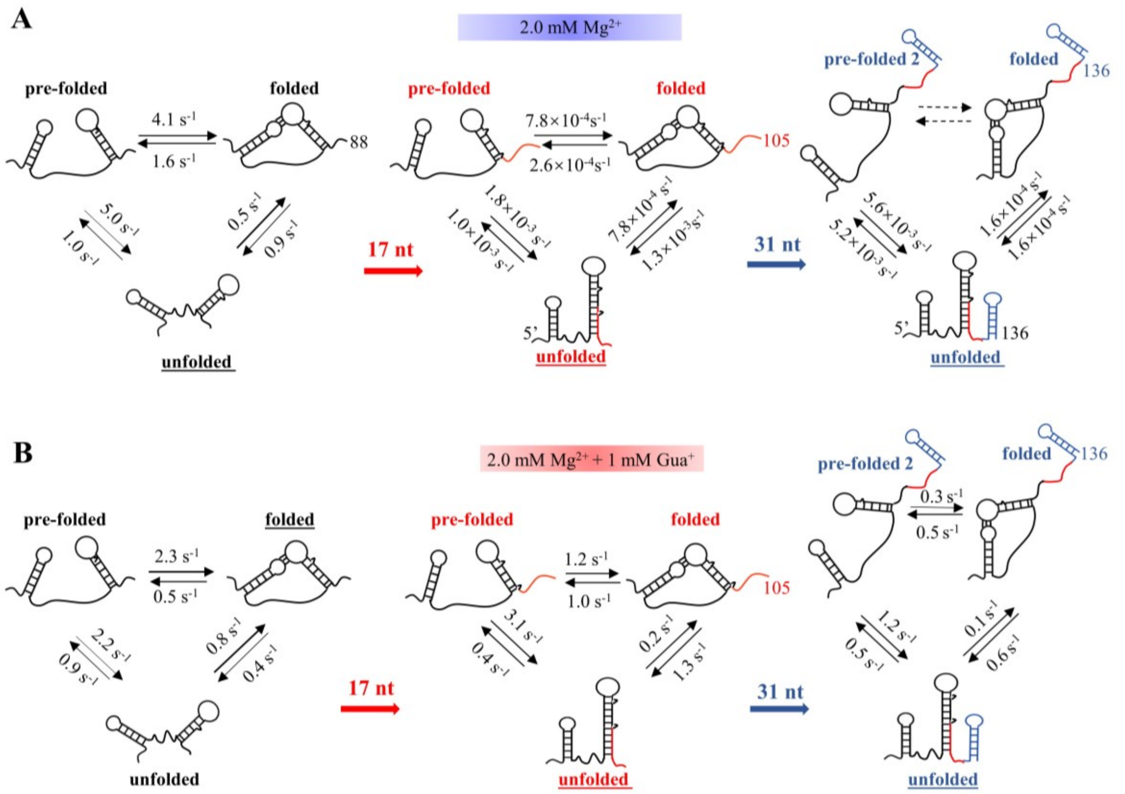


**Figure 5—figure supplement 4 │ Kinetics analysis of the unfolded, pre-folded or pre-folded 2 and folded states of riboG-apt, riboG-term and full-length riboG without and with guanidine at 2 mM $Mg^{2+}$.** The stable structures at each stage are underlined.

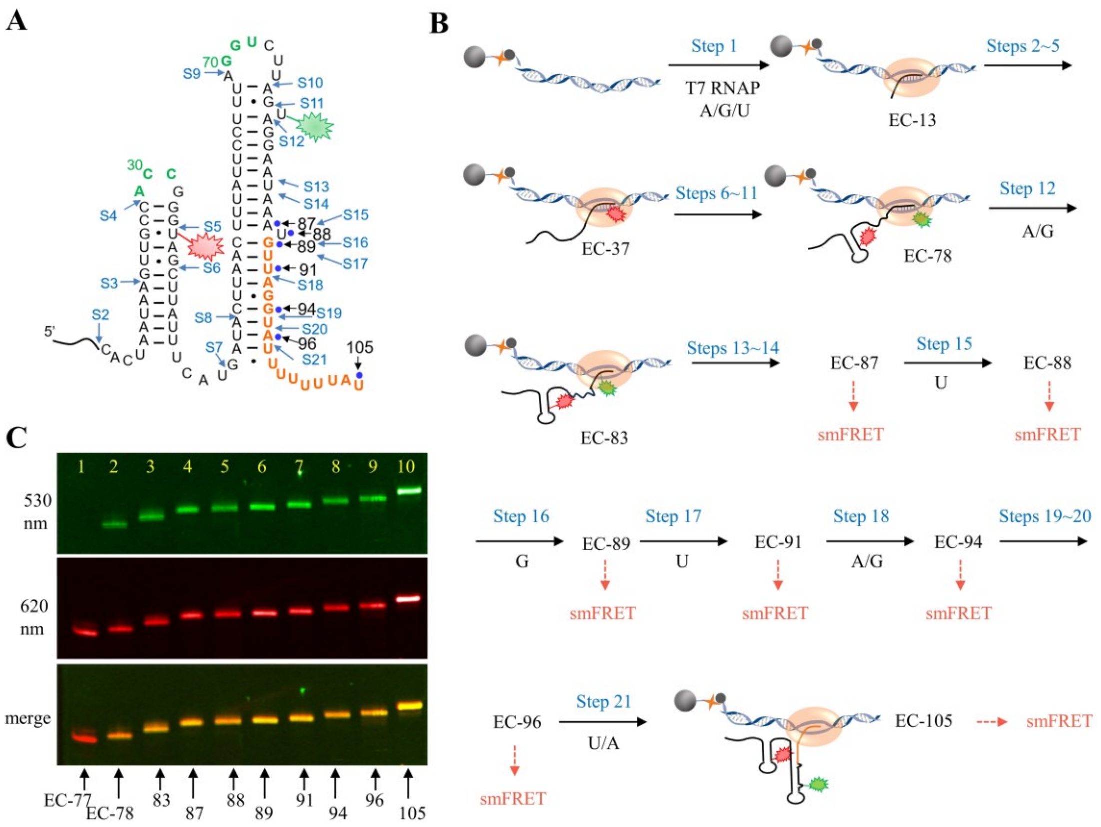


**Figure 6—figure supplement 1 │ The schematic procedure of preparing EC-87 to EC-105 for smFRET study.** A, The secondary structure of riboG-term. The start sites at each step of PLOR were marked by blue arrows. The donor, Cy3 (green sparkle) and acceptor, Cy5 (red sparkle) were localized at U78 and U35, respectively. The paused sites at nascent RNA in ECs were marked with blue dots. B, The schematic procedure of 21-step PLOR reaction for preparing ECs for smFRET study. The reagent usages for the reaction are listed in Supplementary Table S7. C, PAGE images of EC-77 (lane 1), EC-78 (lane 2), EC-83 (lane 3), EC-87 (lane 4), EC-88 (lane 5), EC-89 (lane 6), EC-91 (lane 7), EC-94 (lane 8), EC-96 (lane 9) and EC-105 (lane 10). The top and middle images were irradiated by 530 and 620 nm fluorescence, respectively. The bottom image was merged of the top and middle images.

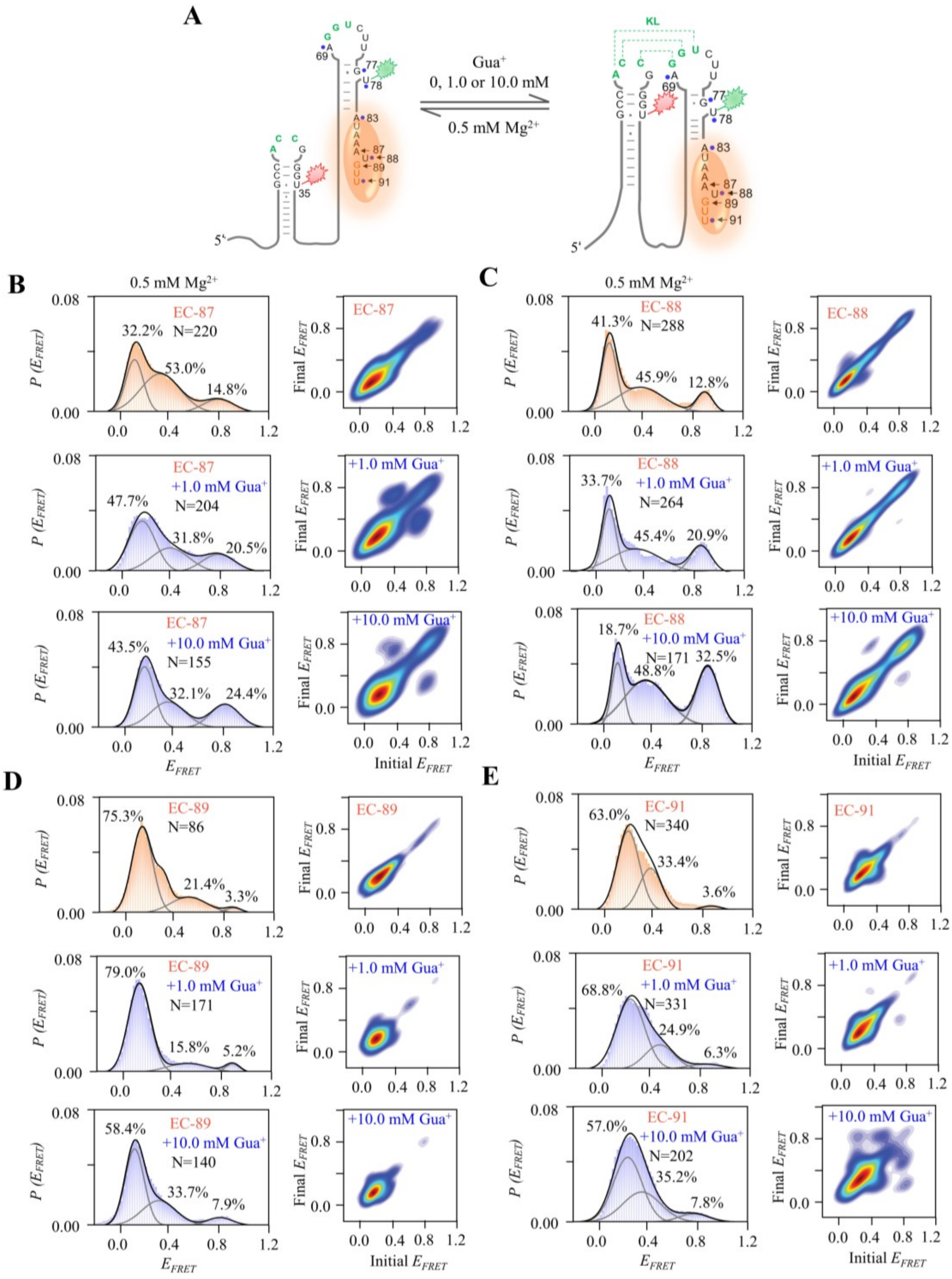

**Figure 6—figure supplement 2 │ smFRET studies of EC-87, EC-88, EC-89, and EC-91 at 0, 1.0 and 10.0 mM Gua$^{+}$ in the presence of 0.5 mM Mg$^{2+}$.** A, The schematic diagram of ECs with nascent RNA and T7 RNA polymerase. The orange ellipse is RNA polymerase. B–E, smFRET histograms and transition density plots of EC-87 (B), EC-88 (C), EC-89 (D), and EC-91(E) at 0 mM, 1.0 mM and 10.0 mM Gua$^{+}$ in the presence of 0.5 mM Mg$^{2+}$.

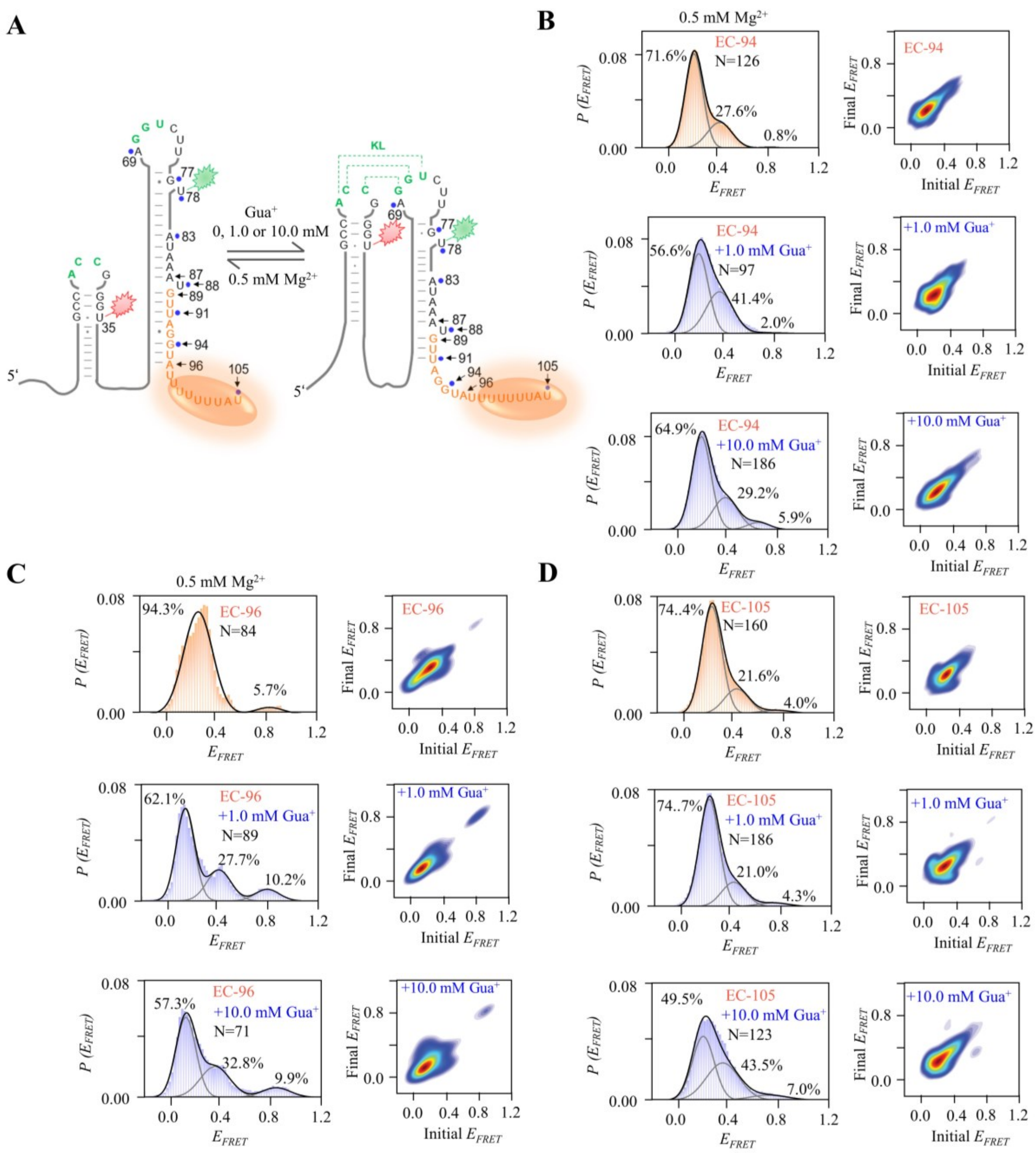

**Figure 6—figure supplement 3 │ smFRET studies of EC-94, EC-96 and EC-105 at 0, 1.0 and 10.0 mM Gua$^+$ in the presence of 0.5 mM Mg$^{2+}$.** A, The schematic diagram of ECs. The orange ellipse is T7 RNA polymerase. B–D, The smFRET histograms and transition density plots of EC-94 (B), EC-96 (C) and EC-105 (D) at 0 mM, 1.0 mM and 10.0 mM Gua$^+$ in the presence of 0.5 mM Mg$^{2+}$.

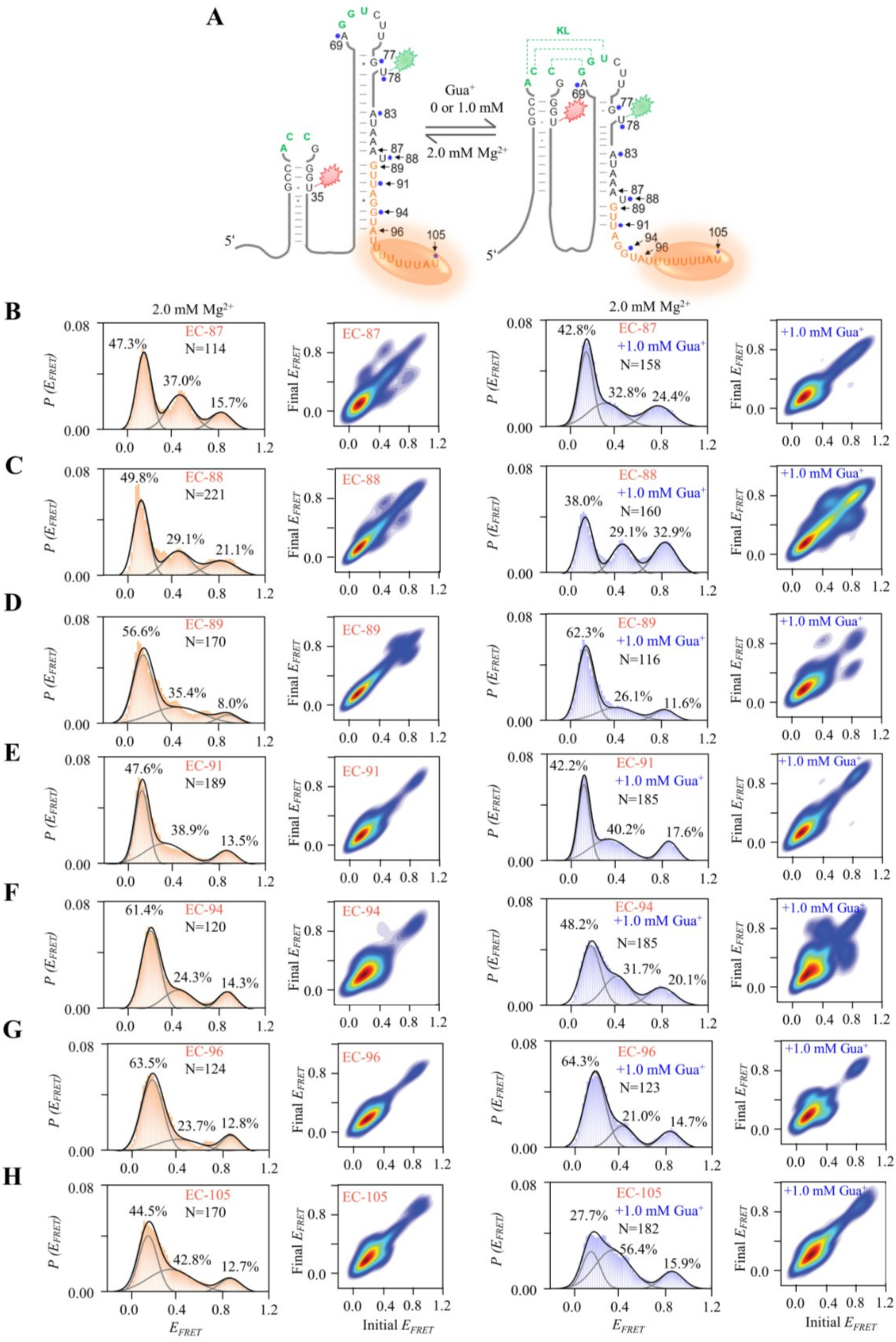


**Figure 6—figure supplement 4 │ smFRET studies of EC-87, EC-88, EC-89, EC-91, EC-94, EC-96 and EC-105 at 0 and 1.0 mM Gua$^{+}$ in the presence of 2.0 mM Mg$^{2+}$.**

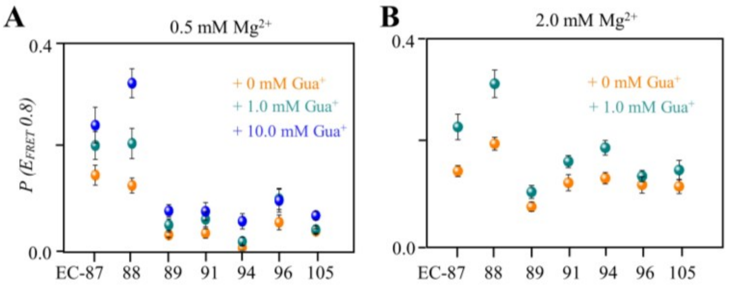


**Figure 6—figure supplement 5 │ Comparison of the folded-conformation percentages in EC-87, EC-88, EC-89, EC-91, EC-94, EC-96 and EC-105 at different $Gua^+$ in the presence of 0.5 mM $Mg^{2+}$ (A) and 2.0 mM $Mg^{2+}$ (B).** The data points of 0 mM, 1.0 mM and 10.0 mM $Gua^+$ are shown in orange, cyan and blue, respectively.

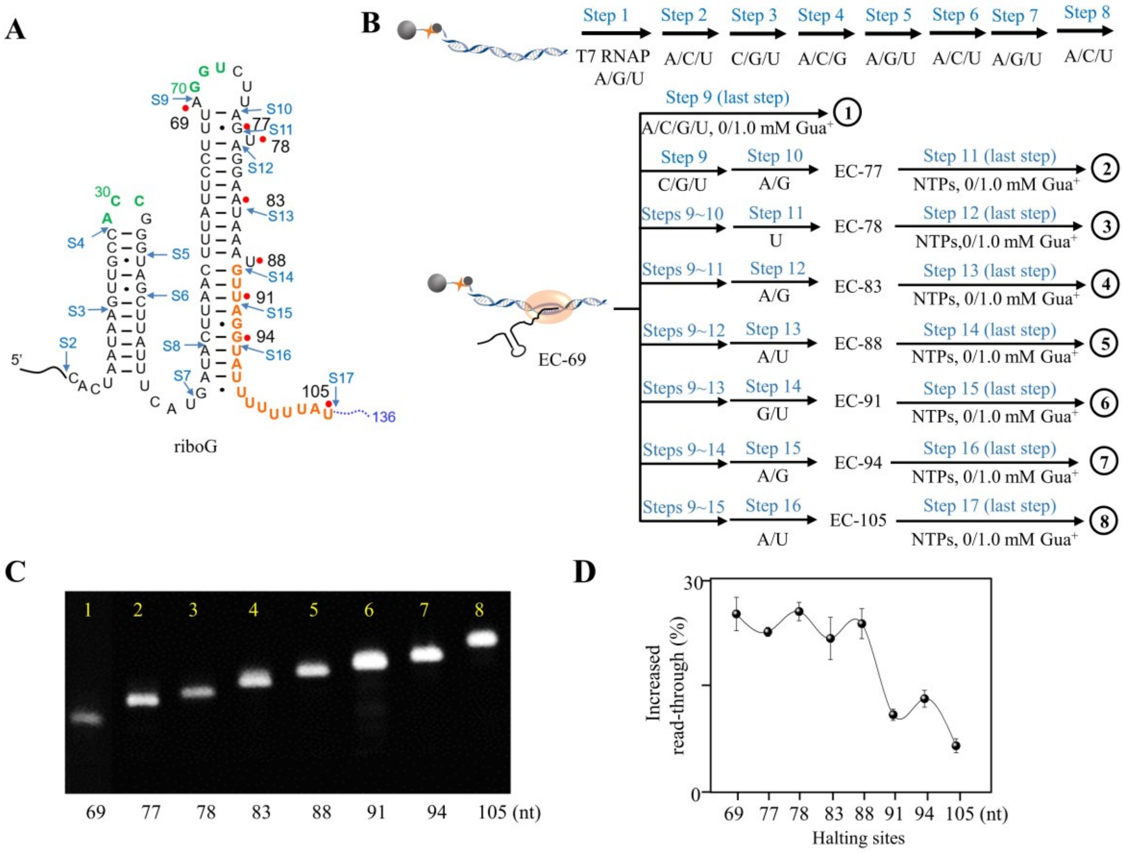


**Figure 7—figure supplement 1 │ *In vitro* transcriptional termination assay of the guanidine-IV riboswitch by PLOR.** A, The full-length guanidine-IV riboswitch can be transcribed by PLOR, halting at specific sites in individual step. The start sites of the 17 step-PLOR reaction were marked by blue arrows. B, Strategies of 9 step- (1), 11 step- (2), 12 step- (3), 13 step- (4), 14 step- (5), 15 step- (6), 16 step- (7) and 17 step-PLOR (8) reactions for single-round transcriptional termination assay. The first 8 steps were shared among different strategies, and 6.0 mM $Mg^{2+}$ was added at each step while 1.0 mM $Gua^{+}$ was added only at the last step. The reagent usages for the 8 reactions are listed in Supplementary Tables S9–16. (C) The denaturing PAGE image of EC-69 to EC-105. The slow migration from EC-69 to EC-105 matches the growing lengths of RNAs. (D) The increase of transcriptional read-through upon the addition of 1 mM $Gua^{+}$ with the last pause site at 69, 77, 78, 83, 88, 91, 94 and 105, respectively before completing transcription.

### Supplement Table S1. The sequences of DNA templates and RNA.

| RNA/DNA | Sequence (5'-3') |
|---|---|
| riboG-apt | GGGAAGAUAUAAUCACUAAUAAGUUGCCACCGGGUAGCUUAUUUCAUGAUACUUAACUUUAUUCCUUUAGGUCUUAGUAGGAAUAAAU |
| riboG-term | GGGAAGAUAUAAUCACUAAUAAGUUGCCACCGGGUAGCUUAUUUCAUGAUACUUAACUUUAUUCCUUUAGGUCUUAGUAGGAAUAAAUGUUAGGUAUUUUUUUAU |
| full-length riboG | GGGAAGAUAUAAUCACUAAUAAGUUGCCACCGGGUAGCUUAUUUCAUGAUACUUAACUUUAUUCCUUUAGGUCUUAGUAGGAAUAAAUGUUAGGUAUUUUUUUAUGUUUAUUUUUAGGAGGAAUCACGAAAAUGAG |
| DNA template for riboG-apt | ATTTATTCCTACTAAGACCTAAAGGAATAAAGTTAAGTATCATGAAATAAGCTACCCGGTGGCAACTTATTAGTGATTATATCTTCCCTATAGTGAGTCGTATTA*TGGACTAGCTGAATCAGA* |
| DNA non-template for riboG-apt | Biotin-*TCTGATTCAGCTAGTCCA*TAATACGACTCACTATAGGGAAGATATAATCACTAATAAGTTGCCACCGGGTAGCTTATTTCATGATACTTAACTTTATTCCTTTAGGTCTTAGTAGGAATAAAT |
| DNA template for riboG-term | ATAAAAAAATACCTAACATTTATTCCTACTAAGACCTAAAGGAATAAAGTTAAGTATCATGAAATAAGCTACCCGGTGGCAACTTATTAGTGATTATATCTTCCCTATAGTGAGTCGTATTA*TGGACTAGCTGAATCAGA* |
| DNA non-template for riboG-term | Biotin-*TCTGATTCAGCTAGTCCA*TAATACGACTCACTATAGGGAAGATATAATCACTAATAAGTTGCCACCGGGTAGCTTATTTCATGATACTTAACTTTATTCCTTTAGGTCTTAGTAGGAATAAATGTTAGGTATTTTTTTAT |
| DNA template for full-length riboG | CTCATTTTCGTGATTCCTCCTAAAAATAAACATAAAAAAATACCTAACATTTATTCCTACTAAGACCTAAAGGAATAAAGTTAAGTATCATGAAATAAGCTACCCGGTGGCAACTTATTAGTGATTATATCTTCCCTATAGTGAGTCGTATTA*TGGACTAGCTGAATCAGA* |
| DNA non-template for full-length riboG | Biotin-*TCTGATTCAGCTAGTCCA*TAATACGACTCACTATAGGGAAGATATAATCACTAATAAGTTGCCACCGGGTAGCTTATTTCATGATACTTAACTTTATTCCTTTAGGTCTTAGTAGGAATAAATGTTAGGTATTTTTTTATGTTTATTTTTAGGAGGAATCACGAAAATGAG |
| Biotinylated DNA | ATTATATCTTCCC-biotin |

The T7 promoter sequences are in blue, and the DNA linker sequences to alleviate steric hindrance are in italic. The underlined sequences in RNA are hybridized with the biotinylated DNA for smFRET.

**Supplement Table S2. The sequences of riboG mutants.**

| RNA/DNA | Sequence (5'-3') |
|---|---|
| riboG-G71C-apt | GGGAAGAUAUAAUCACUAAUAAGUUGCCACCGGGUAGCUUAUUU CAUGAUACUUAACUUUAUUCCUUUAGCUCUUAGUAGGAAUAAAU |
| riboG-G71C-term | GGGAAGAUAUAAUCACUAAUAAGUUGCCACCGGGUAGCUUAUUU CAUGAUACUUAACUUUAUUCCUUUAGCUCUUAGUAGGAAUAAAU GUUAGGUAUUUUUUUAU |
| full-length riboG-G71C | GGGAAGAUAUAAUCACUAAUAAGUUGCCACCGGGUAGCUUAUUU CAUGAUACUUAACUUUAUUCCUUUAGCUCUUAGUAGGAAUAAAU GUUAGGUAUUUUUUUAUGUUUAUUUUUAGGAGGAAUCACGAAAA UGAG |
| riboG-C30G-G71C-apt | GGGAAGAUAUAAUCACUAAUAAGUUGCCAGCGGGUAGCUUAUUU CAUGAUACUUAACUUUAUUCCUUUAGCUCUUAGUAGGAAUAAAU |
| riboG-C30G-G71C-term | GGGAAGAUAUAAUCACUAAUAAGUUGCCAGCGGGUAGCUUAUUU CAUGAUACUUAACUUUAUUCCUUUAGCUCUUAGUAGGAAUAAAU GUUAGGUAUUUUUUUAU |
| full-length riboG-C30G-G71C | GGGAAGAUAUAAUCACUAAUAAGUUGCCAGCGGGUAGCUUAUUU CAUGAUACUUAACUUUAUUCCUUUAGCUCUUAGUAGGAAUAAAU GUUAGGUAUUUUUUUAUGUUUAUUUUUAGGAGGAAUCACGAAAA UGAG |
| riboG-U72C-apt | GGGAAGAUAUAAUCACUAAUAAGUUGCCACCGGGUAGCUUAUUU CAUGAUACUUAACUUUAUUCCUUUAGGCCUUAGUAGGAAUAAAU |
| riboG-U72C-term | GGGAAGAUAUAAUCACUAAUAAGUUGCCACCGGGUAGCUUAUUU CAUGAUACUUAACUUUAUUCCUUUAGGCCUUAGUAGGAAUAAAU GUUAGGUAUUUUUUUAU |
| full-length riboG-U72C | GGGAAGAUAUAAUCACUAAUAAGUUGCCACCGGGUAGCUUAUUU CAUGAUACUUAACUUUAUUCCUUUAGGCCUUAGUAGGAAUAAAU GUUAGGUAUUUUUUUAUGUUUAUUUUUAGGAGGAAUCACGAAAA UGAG |
| riboG-A29G-G72C-apt | GGGAAGAUAUAAUCACUAAUAAGUUGCCGCCGGGUAGCUUAUUU CAUGAUACUUAACUUUAUUCCUUUAGGCCUUAGUAGGAAUAAAU |
| riboG-A29G-G72C-term | GGGAAGAUAUAAUCACUAAUAAGUUGCCGCCGGGUAGCUUAUUU CAUGAUACUUAACUUUAUUCCUUUAGGCCUUAGUAGGAAUAAAU GUUAGGUAUUUUUUUAU |
| full-length riboG-A29G-G72C | GGGAAGAUAUAAUCACUAAUAAGUUGCCGCCGGGUAGCUUAUUU CAUGAUACUUAACUUUAUUCCUUUAGGCCUUAGUAGGAAUAAAU GUUAGGUAUUUUUUUAUGUUUAUUUUUAGGAGGAAUCACGAAAA UGAG |
| riboG-G77C | GGGAAGAUAUAAUCACUAAUAAGUUGCCACCGGGUAGCUUAUUU CAUGAUAGUUAACUUUAUUCCUUUAGGUCUUACUAGGAAUAAAU |

The underlined sequences in RNA are hybridized with the biotinylated DNA for smFRET. The mutated nucleotides are in red.

**Supplement Table S3. Reagent usages for generating Cy3Cy5-riboG-apt.**

| **Reagent usage in a PLOR reaction (100 µL, 10 µM)** |
|---|
| **Step 1** in the buffer (6 mM $MgSO_4$, 40 mM Tris-HCl, 100 mM $K_2SO_4$, 10 mM DTT, pH 8.0) at 37 ℃ for 15 min: 10 µM DNA beads, 10 µM T7 RNAP, 960 µM ATP, 960 µM GTP, 96 µM UTP<br><br>**Steps 2-12** in the buffer (6 mM $MgSO_4$, 40 mM Tris-HCl, 10 mM DTT, pH 8.0) at 25 ℃ for 10 min:<br>**Step 2:** 20 µM CTP, 20 µM UTP, 50 µM ATP<br>**Step 3:** 20 µM GTP, 20 µM UTP, 20 µM CTP<br>**Step 4:** 10 µM ATP, 20 µM CTP, 30 µM GTP<br>**Step 5:** 3 µM Cy5-UTP, 8 µM ATP, 8 µM GTP<br>**Step 6:** 16 µM CTP, 16 µM ATP, 48 µM UTP<br>**Step 7:** 8 µM GTP, 8 µM UTP, 16 µM ATP<br>**Step 8:** 32 µM CTP, 32 µM ATP, 80 µM UTP<br>**Step 9:** 16 µM GTP, 24 µM UTP, 8 µM CTP<br>**Step 10:** 8 µM ATP, 8 µM GTP<br>**Step 11:** 3 µM Cy3-UTP<br>**Step 12:** 16 µM GTP, 16 µM UTP, 48 µM ATP |

**Supplement Table S4. Reagent usages for generating Cy3Cy5-riboG-term.**

| **Reagent usage in a PLOR reaction (100 µL, 10 µM)** |
|---|
| **Step 1** in the buffer (6 mM $MgSO_4$, 40 mM Tris-HCl, 100 mM $K_2SO_4$, 10 mM DTT, pH 8.0) at 37 ℃ for 15 min: 10 µM DNA beads, 10 µM T7 RNAP, 960 µM ATP, 960 µM GTP, 96 µM UTP;<br><br>**Steps 2-13** in the buffer (6 mM $MgSO_4$, 40 mM Tris-HCl, 10 mM DTT, pH 8.0) at 25 ℃ for 10 min:<br>**Step 2:** 20 µM CTP, 20 µM UTP, 50 µM ATP;<br>**Step 3:** 20 µM GTP, 20 µM UTP, 20 µM CTP;<br>**Step 4:** 10 µM ATP, 20 µM CTP, 30 µM GTP;<br>**Step 5:** 3 µM Cy5-UTP, 8 µM ATP, 8 µM GTP;<br>**Step 6:** 16 µM CTP, 16 µM ATP, 48 µM UTP;<br>**Step 7:** 8 µM GTP, 8 µM UTP, 16 µM ATP;<br>**Step 8:** 32 µM CTP, 32 µM ATP, 80 µM UTP;<br>**Step 9:** 16 µM GTP, 24 µM UTP, 8 µM CTP;<br>**Step 10:** 8 µM ATP, 8 µM GTP;<br>**Step 11:** 3 µM Cy3-UTP;<br>**Step 12:** 16 µM GTP, 24 µM ATP;<br>**Step 13:** 24 µM GTP, 48 µM ATP, 104 µM UTP. |

**Supplement Table S5. Reagent usages for generating Cy3Cy5-full-length riboG.**

| **Reagent usage in a PLOR reaction (100 µL, 10 µM)** |
|---|
| **Step 1** in the buffer (6 mM $MgSO_4$, 40 mM Tris-HCl, 100 mM $K_2SO_4$, 10 mM DTT, pH 8.0) at 37 ℃ for 15 min: 10 µM DNA beads, 10 µM T7 RNAP, 960 µM ATP, 960 µM GTP, 96 µM UTP;<br><br>**Steps 2-13** in the buffer (6 mM $MgSO_4$, 40 mM Tris-HCl, 10 mM DTT, pH 8.0) at 25 ℃ for 10 min:<br>**Step 2:** 20 µM CTP, 20 µM UTP, 50 µM ATP;<br>**Step 3:** 20 µM GTP, 20 µM UTP, 20 µM CTP;<br>**Step 4:** 10 µM ATP, 20 µM CTP, 30 µM GTP;<br>**Step 5:** 3 µM Cy5-UTP, 8 µM ATP, 8 µM GTP;<br>**Step 6:** 16 µM CTP, 16 µM ATP, 48 µM UTP;<br>**Step 7:** 8 µM GTP, 8 µM UTP, 16 µM ATP;<br>**Step 8:** 32 µM CTP, 32 µM ATP, 80 µM UTP;<br>**Step 9:** 16 µM GTP, 24 µM UTP, 8 µM CTP;<br>**Step 10:** 8 µM ATP, 8 µM GTP;<br>**Step 11:** 3 µM Cy3-UTP;<br>**Step 12:** 16 µM GTP, 24 µM ATP;<br>**Step 13:** 88 µM GTP, 136 µM ATP, 16 µM CTP, 184 µM UTP. |

### Supplement Table S6. Sequences of RNA in the ECs.

| ECs | Sequences of RNA |
|---|---|
| EC-87 | GGGAAGAUAUAAUCACUAAUAAGUUGCCACCGGGUAGCUU AUUUCAUGAUACUUAACUUUAUUCCUUUAGGUCUUAGUAG GAAUAAA |
| EC-88 | GGGAAGAUAUAAUCACUAAUAAGUUGCCACCGGGUAGCUU AUUUCAUGAUACUUAACUUUAUUCCUUUAGGUCUUAGUAG GAAUAAAU |
| EC-89 | GGGAAGAUAUAAUCACUAAUAAGUUGCCACCGGGUAGCUU AUUUCAUGAUACUUAACUUUAUUCCUUUAGGUCUUAGUAG GAAUAAAUG |
| EC-91 | GGGAAGAUAUAAUCACUAAUAAGUUGCCACCGGGUAGCUU AUUUCAUGAUACUUAACUUUAUUCCUUUAGGUCUUAGUAG GAAUAAAUGUU |
| EC-94 | GGGAAGAUAUAAUCACUAAUAAGUUGCCACCGGGUAGCUU AUUUCAUGAUACUUAACUUUAUUCCUUUAGGUCUUAGUAG GAAUAAAUGUUAGG |
| EC-96 | GGGAAGAUAUAAUCACUAAUAAGUUGCCACCGGGUAGCUU AUUUCAUGAUACUUAACUUUAUUCCUUUAGGUCUUAGUAG GAAUAAAUGUUAGGUA |
| EC-105 | GGGAAGAUAUAAUCACUAAUAAGUUGCCACCGGGUAGCUU AUUUCAUGAUACUUAACUUUAUUCCUUUAGGUCUUAGUAG GAAUAAAUGUUAGGUAUUUUUUUAU |

**Supplement Table S7. Reagent usages for generating EC-87, EC-88, EC-89, EC-91, EC-94, EC-96 and EC-105.**

| **Reagent usage in a PLOR reaction (200 μL, 10 μM)** |
| --- |
| **Step 1** in the buffer (6 mM $MgSO_4$, 40 mM Tris-HCl, 100 mM $K_2SO_4$, 10 mM DTT, pH 8.0) at 37 ℃ for 15 min: 10 μM DNA beads, 10 μM T7 RNAP, 960 μM ATP, 960 μM GTP, 96 μM UTP;<br><br>**Steps 2-21** in the buffer (6 mM $MgSO_4$, 40 mM Tris-HCl, 10 mM DTT, pH 8.0) at 25 ℃ for 10 min:<br>**Step 2:** 20 μM CTP, 20 μM UTP, 50 μM ATP;<br>**Step 3:** 20 μM GTP, 20 μM UTP, 20 μM CTP;<br>**Step 4:** 10 μM ATP, 20 μM CTP, 30 μM GTP;<br>**Step 5:** 3 μM Cy5-UTP, 8 μM ATP, 8 μM GTP;<br>**Step 6:** 16 μM CTP, 16 μM ATP, 48 μM UTP;<br>**Step 7:** 8 μM GTP, 8 μM UTP, 16 μM ATP;<br>**Step 8:** 32 μM CTP, 32 μM ATP, 80 μM UTP;<br>**Step 9:** 16 μM GTP, 24 μM UTP, 8 μM CTP;<br>**Step 10:** 8 μM ATP, 8 μM GTP;<br>**Step 11:** 3 μM Cy3-UTP;<br>**Step 12:** 16 μM GTP, 24 μM ATP;<br>**Step 13:** 8 μM UTP;<br>**Step 14:** 24 μM ATP (the dissociated EC-87 was used for smFRET);<br>**Step 15:** 8 μM UTP (the dissociated EC-88 was used for smFRET);<br>**Step 16:** 8 μM GTP (the dissociated EC-89 was used for smFRET);<br>**Step 17:** 16 μM UTP (the dissociated EC-91 was used for smFRET);<br>**Step 18:** 8 μM ATP, 16 μM GTP (the dissociated EC-94 was used for smFRET);<br>**Step 19:** 8 μM UTP;<br>**Step 20:** 8 μM ATP (the dissociated EC-96 was used for smFRET);<br>**Step 21:** 64 μM UTP, 8 μM ATP (the dissociated EC-105 was used for smFRET). |

**Supplement Table S8. Reagent usages for termination assays by 8 step-PLOR.**

| **Reagent usage in PLOR reactions (100 µL, 3 µM)** |
|---|
| **Step 1** in the buffer (6 mM $MgSO_4$, 40 mM Tris-HCl, 100 mM $K_2SO_4$, 10 mM DTT, pH 8.0) at 37 ℃ for 15 min: 3 µM DNA beads, 3 µM T7 RNAP, 288 µM ATP, 288 µM GTP, 28.8 µM UTP; |
| **Steps 2-8** in the buffer (40 mM Tris-HCl, 10 mM DTT, pH 8.0) at 25 ℃ for 10 min:<br>**Step 2:** 6 µM CTP, 6 µM UTP, 15 µM ATP, 6 mM $MgSO_4$;<br>**Step 3:** 6 µM GTP, 6 µM UTP, 6 µM CTP, 6 mM $MgSO_4$;<br>**Step 4:** 3 µM ATP, 6 µM CTP, 9 µM GTP, 6 mM $MgSO_4$;<br>**Step 5:** 0.9 µM UTP, 2.4µM ATP, 2.4µM GTP, 6 mM $MgSO_4$;<br>**Step 6:** 4.8 µM CTP, 4.8 µM ATP, 14.4 µM UTP, 6 mM $MgSO_4$;<br>**Step 7:** 2.4 µM GTP, 2.4 µM UTP, 4.8 µM ATP, 6 mM $MgSO_4$;<br>**Step 8-a:** 6.3 µM CTP, 22.5 µM ATP, 33.3 µM UTP, 14.4 µM GTP, 6 mM $MgSO_4$, 0–10 mM $Gua^+$;<br>**Step 8-b:** 6.3 µM CTP, 22.5 µM ATP, 33.3 µM UTP, 14.4 µM GTP, 2 mM $MgSO_4$, 0–10 mM $Gua^+$. |

**Supplement Table S9. Reagent usages for termination assay by 9 step-PLOR.**

| **Reagent usage in PLOR reactions (100 µL, 3 µM)** |
|---|
| **Step 1** in the buffer (6 mM $MgSO_4$, 40 mM Tris-HCl, 100 mM $K_2SO_4$, 10 mM DTT, pH 8.0) at 37 ℃ for 15 min: 3µM DNA beads, 3 µM T7 RNAP, 288 µM ATP, 288 µM GTP, 28.8 µM UTP; |
| **Steps 2-9** in the buffer (6 mM $MgSO_4$, 40 mM Tris-HCl, 10 mM DTT, pH 8.0) at 25 ℃ for 10 min:<br>**Step 2:** 6 µM CTP, 6 µM UTP, 15 µM ATP;<br>**Step 3:** 6 µM GTP, 6 µM UTP, 6 µM CTP;<br>**Step 4:** 3 µM ATP, 6 µM CTP, 9 µM GTP;<br>**Step 5:** 0.9 µM UTP, 2.4µM ATP, 2.4µM GTP;<br>**Step 6:** 4.8 µM CTP, 4.8 µM ATP, 14.4 µM UTP;<br>**Step 7:** 2.4 µM GTP, 2.4 µM UTP, 4.8 µM ATP;<br>**Step 8:** 9.6 µM CTP, 24 µM UTP, 9.6 µM ATP;<br>**Step 9:** 2.7 µM CTP, 18.9 µM ATP, 24.3 µM UTP, 14.4 µM GTP, 0 or 1 mM $Gua^{+}$. |

**Supplement Table S10. Reagent usages for termination assay by 11 step-PLOR.**

| **Reagent usage in PLOR reactions (100 µL, 3 µM)** |
|---|
| **Step 1** in the buffer (6 mM $MgSO_4$, 40 mM Tris-HCl, 100 mM $K_2SO_4$, 10 mM DTT, pH 8.0) at 37 ℃ for 15 min: 3 µM DNA beads, 3 µM T7 RNAP, 288 µM ATP, 288 µM GTP, 28.8 µM UTP; |
| **Steps 2-11** in the buffer (6 mM $MgSO_4$, 40 mM Tris-HCl, 10 mM DTT, pH 8.0) at 25 ℃ for 10 min:<br>**Step 2:** 6 µM CTP, 6 µM UTP, 15 µM ATP;<br>**Step 3:** 6 µM GTP, 6 µM UTP, 6 µM CTP;<br>**Step 4:** 3 µM ATP, 6 µM CTP, 9 µM GTP;<br>**Step 5:** 0.9 µM UTP, 2.4µM ATP, 2.4µM GTP;<br>**Step 6:** 4.8 µM CTP, 4.8 µM ATP, 14.4 µM UTP;<br>**Step 7:** 2.4 µM GTP, 2.4 µM UTP, 4.8 µM ATP;<br>**Step 8:** 9.6 µM CTP, 24 µM UTP, 9.6 µM ATP;<br>**Step 9:** 4.8 µM GTP, 7.2 µM UTP, 2.4 µM CTP;<br>**Step 10:** 2.4 µM GTP, 2.4 µM ATP;<br>**Step 11:** 1.8 µM CTP, 18 µM ATP, 21.6 µM UTP, 11.7 µM GTP, 0 or 1 mM $Gua^+$. |

**Supplement Table S11. Reagent usages for termination assay by 12 step-PLOR.**

| **Reagent usage in PLOR reactions (100 μL, 3 μM)** |
|---|
| **Step 1** in the buffer (6 mM $MgSO_4$, 40 mM Tris-HCl, 100 mM $K_2SO_4$, 10 mM DTT, pH 8.0) at 37 ℃ for 15 min: 3 μM DNA beads, 3 μM T7 RNAP, 288 μM ATP, 288 μM GTP, 28.8 μM UTP;<br><br>**Steps 2-12** in the buffer (6 mM $MgSO_4$, 40 mM Tris-HCl, 10 mM DTT, pH 8.0) at 25 ℃ for 10 min:<br>**Step 2:** 6 μM CTP, 6 μM UTP, 15 μM ATP;<br>**Step 3:** 6 μM GTP, 6 μM UTP, 6 μM CTP;<br>**Step 4:** 3 μM ATP, 6 μM CTP, 9 μM GTP;<br>**Step 5:** 0.9 μM UTP, 2.4μM ATP, 2.4μM GTP;<br>**Step 6:** 4.8 μM CTP, 4.8 μM ATP, 14.4 μM UTP;<br>**Step 7:** 2.4 μM GTP, 2.4 μM UTP, 4.8 μM ATP;<br>**Step 8:** 9.6 μM CTP, 24 μM UTP, 9.6 μM ATP;<br>**Step 9:** 4.8 μM GTP, 7.2 μM UTP, 2.4 μM CTP;<br>**Step 10:** 2.4 μM GTP, 2.4 μM ATP;<br>**Step 11:** 2.4 μM UTP;<br>**Step 12:** 1.8 μM CTP, 18 μM ATP, 20.7 μM UTP, 11.7 μM GTP, 0 or 1 mM $Gua^+$. |

**Supplement Table S12. Reagent usages for termination assay by 13 step-PLOR.**

| **Reagent usage in PLOR reactions (100 μL, 3 μM)** |
|---|
| **Step 1** in the buffer (6 mM $MgSO_4$, 40 mM Tris-HCl, 100 mM $K_2SO_4$, 10 mM DTT, pH 8.0) at 37 ℃ for 15 min: 3 μM DNA beads, 3 μM T7 RNAP, 288 μM ATP, 288 μM GTP, 28.8 μM UTP;<br><br>**Steps 2-13** in the buffer (6 mM $MgSO_4$, 40 mM Tris-HCl, 10 mM DTT, pH 8.0) at 25 ℃ for 10 min:<br>**Step 2:** 6 μM CTP, 6 μM UTP, 15 μM ATP;<br>**Step 3:** 6 μM GTP, 6 μM UTP, 6 μM CTP;<br>**Step 4:** 3 μM ATP, 6 μM CTP, 9 μM GTP;<br>**Step 5:** 0.9 μM UTP, 2.4μM ATP, 2.4μM GTP;<br>**Step 6:** 4.8 μM CTP, 4.8 μM ATP, 14.4 μM UTP;<br>**Step 7:** 2.4 μM GTP, 2.4 μM UTP, 4.8 μM ATP;<br>**Step 8:** 9.6 μM CTP, 24 μM UTP, 9.6 μM ATP;<br>**Step 9:** 4.8 μM GTP, 7.2 μM UTP, 2.4 μM CTP;<br>**Step 10:** 2.4 μM GTP, 2.4 μM ATP;<br>**Step 11:** 2.4 μM UTP;<br>**Step 12:** 4.8 μM GTP, 7.2 μM ATP;<br>**Step 13:** 1.8 μM CTP, 15.3 μM ATP, 20.7 μM UTP, 9.9 μM GTP, 0 or 1 mM $Gua^+$. |

**Supplement Table S13. Reagent usages for termination assay by 14 step-PLOR.**

| **Reagent usage in PLOR reactions (100 µL, 3 µM)** |
|---|
| **Step 1** in the buffer (6 mM $MgSO_4$, 40 mM Tris-HCl, 100 mM $K_2SO_4$, 10 mM DTT, pH 8.0) at 37 ℃ for 15 min: 3 µM DNA beads, 3 µM T7 RNAP, 288 µM ATP, 288 µM GTP, 28.8 µM UTP;<br><br>**Steps 2-14** in the buffer (6 mM $MgSO_4$, 40 mM Tris-HCl, 10 mM DTT, pH 8.0) at 25 ℃ for 10 min:<br>**Step 2:** 6 µM CTP, 6 µM UTP, 15 µM ATP;<br>**Step 3:** 6 µM GTP, 6 µM UTP, 6 µM CTP;<br>**Step 4:** 3 µM ATP, 6 µM CTP, 9 µM GTP;<br>**Step 5:** 0.9 µM UTP, 2.4µM ATP, 2.4µM GTP;<br>**Step 6:** 4.8 µM CTP, 4.8 µM ATP, 14.4 µM UTP;<br>**Step 7:** 2.4 µM GTP, 2.4 µM UTP, 4.8 µM ATP;<br>**Step 8:** 9.6 µM CTP, 24 µM UTP, 9.6 µM ATP;<br>**Step 9:** 4.8 µM GTP, 7.2 µM UTP, 2.4 µM CTP;<br>**Step 10:** 2.4 µM GTP, 2.4 µM ATP;<br>**Step 11:** 2.4 µM UTP;<br>**Step 12:** 4.8 µM GTP, 7.2 µM ATP;<br>**Step 13:** 4.8 µM UTP, 7.2 µM ATP;<br>**Step 14:** 1.8 µM CTP, 12.6 µM ATP, 18.9 µM UTP, 9.9 µM GTP, 0 or 1 mM $Gua^+$. |

**Supplement Table S14. Reagent usages for termination assay by 15 step-PLOR.**

| **Reagent usage in PLOR reactions (100 μL, 3 μM)** |
|---|
| **Step 1** in the buffer (6 mM $MgSO_4$, 40 mM Tris-HCl, 100 mM $K_2SO_4$, 10 mM DTT, pH 8.0) at 37 ℃ for 15 min: 3 μM DNA beads, 3 μM T7 RNAP, 288 μM ATP, 288 μM GTP, 28.8 μM UTP;<br><br>**Steps 2-15** in the buffer (6 mM $MgSO_4$, 40 mM Tris-HCl, 10 mM DTT, pH 8.0) at 25 ℃ for 10 min:<br>**Step 2:** 6 μM CTP, 6 μM UTP, 15 μM ATP;<br>**Step 3:** 6 μM GTP, 6 μM UTP, 6 μM CTP;<br>**Step 4:** 3 μM ATP, 6 μM CTP, 9 μM GTP;<br>**Step 5:** 0.9 μM UTP, 2.4μM ATP, 2.4μM GTP;<br>**Step 6:** 4.8 μM CTP, 4.8 μM ATP, 14.4 μM UTP;<br>**Step 7:** 2.4 μM GTP, 2.4 μM UTP, 4.8 μM ATP;<br>**Step 8:** 9.6 μM CTP, 24 μM UTP, 9.6 μM ATP;<br>**Step 9:** 4.8 μM GTP, 7.2 μM UTP, 2.4 μM CTP;<br>**Step 10:** 2.4 μM GTP, 2.4 μM ATP;<br>**Step 11:** 2.4 μM UTP;<br>**Step 12:** 4.8 μM GTP, 7.2 μM ATP;<br>**Step 13:** 4.8 μM UTP, 7.2 μM ATP;<br>**Step 14:** 2.4 μM GTP, 4.8 μM UTP;<br>**Step 15:** 1.8 μM CTP, 12.6 μM ATP, 17.1 μM UTP, 9.0 μM GTP, 0 or 1 mM $Gua^+$. |

**Supplement Table S15. Reagent usages for termination assay by 16 step-PLOR.**

| **Reagent usage in PLOR reactions (100 µL, 3 µM)** |
|---|
| **Step 1** in the buffer (6 mM $MgSO_4$, 40 mM Tris-HCl, 100 mM $K_2SO_4$, 10 mM DTT, pH 8.0) at 37 ℃ for 15 min: 3 µM DNA beads, 3 µM T7 RNAP, 288 µM ATP, 288 µM GTP, 28.8 µM UTP;<br><br>**Steps 2-16** in the buffer (6 mM $MgSO_4$, 40 mM Tris-HCl, 10 mM DTT, pH 8.0) at 25 ℃ for 10 min:<br>**Step 2:** 6 µM CTP, 6 µM UTP, 15 µM ATP;<br>**Step 3:** 6 µM GTP, 6 µM UTP, 6 µM CTP;<br>**Step 4:** 3 µM ATP, 6 µM CTP, 9 µM GTP;<br>**Step 5:** 0.9 µM UTP, 2.4µM ATP, 2.4µM GTP;<br>**Step 6:** 4.8 µM CTP, 4.8 µM ATP, 14.4 µM UTP;<br>**Step 7:** 2.4 µM GTP, 2.4 µM UTP, 4.8 µM ATP;<br>**Step 8:** 9.6 µM CTP, 24 µM UTP, 9.6 µM ATP;<br>**Step 9:** 4.8 µM GTP, 7.2 µM UTP, 2.4 µM CTP;<br>**Step 10:** 2.4 µM GTP, 2.4 µM ATP;<br>**Step 11:** 2.4 µM UTP;<br>**Step 12:** 4.8 µM GTP, 7.2 µM ATP;<br>**Step 13:** 4.8 µM UTP, 7.2 µM ATP;<br>**Step 14:** 2.4 µM GTP, 4.8 µM UTP;<br>**Step 15:** 4.8 µM GTP, 2.4 µM ATP;<br>**Step 16:** 1.8 µM CTP, 11.7 µM ATP, 17.1 µM UTP, 7.2 µM GTP, 0 or 1 mM $Gua^+$. |

**Supplement Table S16. Reagent usages for termination assay by 17 step-PLOR.**

| **Reagent usage in PLOR reactions (100 µL, 3 µM)** |
|---|
| **Step 1** in the buffer (6 mM $MgSO_4$, 40 mM Tris-HCl, 100 mM $K_2SO_4$, 10 mM DTT, pH 8.0) at 37 ℃ for 15 min: 3 µM DNA beads, 3 µM T7 RNAP, 288 µM ATP, 288 µM GTP, 28.8 µM UTP;<br><br>**Steps 2-17** in the buffer (6 mM $MgSO_4$, 40 mM Tris-HCl, 10 mM DTT, pH 8.0) at 25 ℃ for 10 min:<br>**Step 2:** 6 µM CTP, 6 µM UTP, 15 µM ATP;<br>**Step 3:** 6 µM GTP, 6 µM UTP, 6 µM CTP;<br>**Step 4:** 3 µM ATP, 6 µM CTP, 9 µM GTP;<br>**Step 5:** 0.9 µM UTP, 2.4µM ATP, 2.4µM GTP;<br>**Step 6:** 4.8 µM CTP, 4.8 µM ATP, 14.4 µM UTP;<br>**Step 7:** 2.4 µM GTP, 2.4 µM UTP, 4.8 µM ATP;<br>**Step 8:** 9.6 µM CTP, 24 µM UTP, 9.6 µM ATP;<br>**Step 9:** 4.8 µM GTP, 7.2 µM UTP, 2.4 µM CTP;<br>**Step 10:** 2.4 µM GTP, 2.4 µM ATP;<br>**Step 11:** 2.4 µM UTP;<br>**Step 12:** 4.8 µM GTP, 7.2 µM ATP;<br>**Step 13:** 4.8 µM UTP, 7.2 µM ATP;<br>**Step 14:** 2.4 µM GTP, 4.8 µM UTP;<br>**Step 15:** 4.8 µM GTP, 2.4 µM ATP;<br>**Step 16:** 21.6 µM UTP, 4.8 µM ATP;<br>**Step 17:** 1.8 µM CTP, 9.9 µM ATP, 9.0 µM UTP, 7.2 µM GTP, 0 or 1 mM $Gua^+$. |
